# Efficient Prediction of Microplastic Counts from Mass Measurements

**DOI:** 10.1101/2021.01.04.425187

**Authors:** Shuyao Tan, Joshua Taylor, Elodie Passeport

## Abstract

Microplastics must be characterized and quantified to assess their impact. Current quantification procedures are time-consuming and prone to human error. This study evaluates the use of machine learning to estimate the number of microplastic particles based on aggregate particle weight measurements. Synthetic datasets are used to test the performance of linear regression, kernel ridge regression and decision trees. Kernel ridge regression achieves the strongest performance, and it is also tested with experimental datasets. The numerical results show that the algorithm is better at predicting the counts of larger and more homogeneous samples, and that contamination by organics does not significantly increase error. In mixed samples, prediction error is lower for heavier particles, with an error rate comparable to or better than that of manual counting. Overall, the proposed method is faster and easier than current approaches.

## 1. Introduction

Plastic particles less than 5 mm in size are referred to as microplastics. ^1^ They are widespread in the environment, both in densely populated and remote areas. ^2-5^ Much remains unknown about their impact on organisms; while negative effects have been observed in several studies, a clear, causal picture has not yet emerged. ^6^ This is in part due to the diversity of microplastics, i.e., shape, size, polymer identity, chemical mixture, color, and count, which are not accounted for in most studies on their ecological effects. ^6,7^

Research on microplastics can have several different objectives, e.g., source identification, fate assessment, and eco)toxicological impact evaluation, all of which require the characterization and quantification of microplastics. Depending on the subject, the quantity of microplastics can be referred to as count, concentration, abundance, or dose, expressed as particles per sampling area, per volume or per mass of sample. ^8–10^ Here, we will use the term count. The count of microplastics, though not sufficient alone to address all research questions, remains an important metric. Indeed, higher concentrations of microplastics are found near more populated areas, thus helping estimate proximity to sources. Microplastic counts are also used to evaluate the performance of water treatment systems. ^11,12^ Finally, the count of microplastics is also one of the drivers of observed effects on organisms. ^6^

While currently there are no formal standards for microplastic characterization and quantification, the most common and non-destructive method involves some or all of the following steps: organic digestion, in which chemicals or enzymes are used to remove organics; density separation, in which a solution of known density is used to separate lighter particles from heavier sediments; sieving, in which the particles are passed through sieves of different mesh sizes and grouped into different size ranges; visual identification, in which particles are manually sorted under a microscope for counting, sometimes particle size measurement, and classification for colors and shape categories. ^8–10,13–15^ The counting step is time-consuming and subject to human error. Usually a technique such as Raman or Fourier-transform infrared spectroscopy FTIR) is also used to verify that the particles are indeed plastic and identify the polymer type, further increasing cost and time. As a result, characterization and quantification have become the rate-limiting steps in the analysis of microplastic contamination.

Recent efforts have attempted to make this process cheaper and faster by automation. Algorithms have been developed to automatically match *μ*FTIR spectra to polymers, which take significantly less time than manual labour and involve less error ^3,16–18^. Various machine learning algorithms have also been coupled with spectroscopic techniques to auto-identify microplastic particles. ^19^ For example, a convolutional neural network was trained to classify microbeads based on microscopic images, and support vector machines, adaptive boosting and random forests have been trained to identify particles from scanned images. ^20,21^ These methods are highly effective, but their applicability is limited for one or more of the following reasons: (i) they rely on expensive equipment, which require expertise to use, (ii) cannot reliably distinguish plastics from organic particles, and (iii) require the particles to be loosely distributed over a surface. Our objective is to develop an alternative that is cheap, fast, and simple enough that anyone can use it to reliably quantify microplastic contamination.

The objective of this study is to use machine learning to estimate the number of microplastic particles, in different size ranges and morphology categories, from their aggregate weight measurements. Microplastic samples will be sieved into different size ranges, and the total particle weight of each size range will be measured. From the resulting set of weight measurements, we estimate the count of microplastic particles in each of the following types: fragment, bead, film, fiber, and rubber. Koelmans et al. have proposed empirical relationships to calculate a microplastic mass-volume-number conversion factor based on data averaged over many microplastic studies. ^22^ In this study, we aim to develop a method for quantifying microplastic particles without assuming that all samples share similar compositions. We use three machine learning algorithms: linear regression, kernel ridge regression, and decision trees. We have chosen these techniques for their simplicity and low computational cost. While neural networks could potentially also work well, we prefer these techniques for several reasons. First, the effectiveness of modern, deep neural networks comes from massive training datasets, which are not available in this application. Second, neural networks are more computationally intensive and often entail extensive parameter tuning, thus increasing the complexity of the procedure. In our numerical results, we will compare the performance of the algorithms to manual counting and evaluate the impact of mass measurement error on count prediction.

## 2. Methods

### 2.1. Experimental and Synthetic Datasets

Five sets of experimental data are taken from the literature. Isobe *et al*.^23^ made samples by adding microplastic particles to sea water, and sent them to different laboratories around the world. These samples were sieved into multiple size ranges, weighed, and counted, resulting in 10 pairs of particle weight and count measurements. McIlwraith *et al*.^24^ washed blankets in a washing machine, collected fibers from the effluent water, and reported 18 pairs of total weight and count. Lavers *et al*.^25^ reported 11 pairs of particle weight and count measurements obtained from sand samples collected in the Cocos islands. Puskic *et al*.^26^ analysed microplastic particles in 52 sea birds (Short-Tailed Shearwaters), resulting in 30 weight and count pairs. Corcoran and Arturo *et al*.^27,28^ collected visible polymeric debris from 66 beaches across the Laurentian Great Lakes, measured the total weight of microplastic particles ranging from 1 mm to 5 mm, and reported particle count by morphology. These datasets will be referred to as *Interlab, Washing Machine, Cocos, Shearwaters*, and *Great Lakes*, respectively.

Due to the limited quantity and variation of published experimental data, a computational procedure was developed to generate synthetic data. These synthetic datasets were used to evaluate the performance of the three algorithms under various circumstances. The procedure is summarized in Table S1. A particle’s density depends on its polymer identity and weathering state. The morphology (hereafter referred to as type) of a particle is defined by its largest, medium, and smallest dimensions. The number of each type of particles in each of the samples described below was randomly drawn from the ranges given in Table S1 in Supporting Information. On average, there can be around 10 rubbers, 40 fragments, 50 films, 50 beads and 300 fibers in a sample. These values are approximates to analysed samples collected from freshwater systems that are subject to human activities. ^3,29–31^ The probability density plots of the number of each type of particle in the synthetic datasets can be found in Section S1 in Supporting Information. The dimensions of each particle were set as follows.

1. Rubber particles and fragments had similar dimensions but different densities. Their largest dimension was uniformly distributed between 153 *μ*m and 1500 *μ*m. The medium dimension was 70 to 100% of the largest dimension. The smallest dimension was 40 to 100% of the medium dimension. Rubber particles were lighter than fragments.
2. Films had similar densities to fragments, but their smallest dimensions were smaller. The largest and medium dimensions of films were drawn from the same range as those of fragments. The smallest dimension for films was 5 to 10% of the medium dimension.
3. Beads were assumed to be perfect spheres. Each bead’s diameter was a uniformly drawn random number between 107 *μ*m and 1500 *μ*m.
4. Fibers were randomly divided into long and short fibers. Their diameters were between 10 and 40 *μ*m. The length of each long fiber was 30 to 50 times of its diameter, and the length of each short fiber was 10 to 30 times of its diameter.
5. The largest dimension of organic particles was between 214 and 2000 *μ*m. The medium dimension was 50 to 100% of the largest dimension. The smallest dimension was between 40 and 150 *μ*m.

The sieving process for non-fiber particles was simulated by comparing the size of each particle with the mesh size. If both the largest and medium dimensions were larger than the mesh size, the particle remained on the sieve; if not, the particle passed through the sieve to the next one. In this study, the sieve mesh sizes were: 1000 *μ*m, 500 *μ*m, 300 *μ*m and 106 *μ*m. Although fibers were thin enough to pass through every sieve, they could still become entangled with the mesh or with other particles. To approximate this randomness, the probability for long fibers to stay on each sieve was set to be 0.3, 0.4, 0.2, 0.1, and for short fibers, the distribution was 0.15, 0.35, 0.3, 0.2. These probabilities were estimated from storm water sample data collected from a parking lot using the same four sieve sizes specified above. ^29^

After sieving, the total mass of all particles on each sieve was calculated by summing up all individual particle masses. Individual masses were obtained by multiplying the volume and density of each particle. For cuboid-shaped particles, volume was calculated by multiplying the three dimensions. Fiber volume was the product of its cross-sectional area and length. Spherical particles were treated as perfect spheres.

In this study, 10 datasets with different particle combinations were generated, each containing 100 samples (Section S2). These 10 combinations were:

1. Film and fiber.
2. Bead and fiber.
3. Rubber, fragment and fiber.
4. Rubber, fragment, fiber and half as many organic particles as fragments.
5. Rubber, fragment, fiber and as many organic particles as fragments.
6. Rubber, fragment, fiber and film.
7. Rubber, fragment, fiber, film and as many organic particles as fragments.
8. Rubber, fragment, fiber and bead.
9. Rubber, fragment, fiber, film and bead.
10. Rubber, fragment, fiber, film, bead and as many organic particles as fragments.

Fibers appeared in all samples because they are the most common type of particle found in water, sediment and air. ^29,32,33^ Fragments appeared in most samples because they are the second most common particle in water and sediment. ^32^ The above synthetic datasets form the base case for the numerical experiments in later sections. To test the dependence of algorithm performance on sample size, two more cases were generated with composition 9, but with more particles, as described in Table S2.

### 2.2. Machine Learning Algorithms

Let *x*_*i*_ = [*x*_*i*1_, *x*_*i*2_, …, *x*_*ik*_] be a row vector of the masses in each of *k* sieves for the *i*^th^sample. The matrix *X* = [*x*_1_; *x*_2_; *x*_3_; …; *x*_*n*_] contains the mass measurements for all *n* samples. Let *y*_*i*_ = [*y*_*i*1_, *y*_*i*2_, …, *y*_*im*_] be a row vector of the number of each type of particle in each sieve for the *i*^th^sample, and *Y* = [*y*_1_; *y*_2_; *y*_3_; …; *y*_*n*_] be the matrix of all counts. Let *Ŷ* = *f* (*X*) be an estimate of *Y* based on some function of *X* In each of the following learning frameworks, we assumed some simple structure for *f*, and then used training data consisting of known (*X, Y*) pairs to solve for the parameters of *f* that minimize prediction error

#### Linear Regression (LR)

Linear regression assumes a relationship of the form

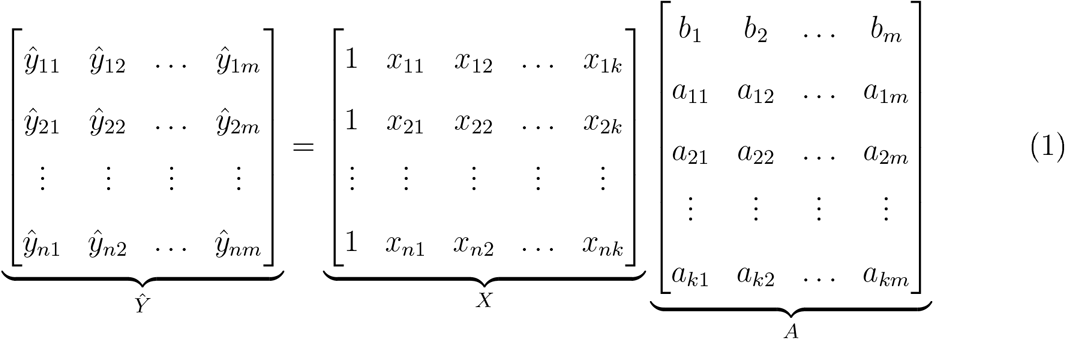

Given training data (*X, Y*), the matrix A that minimizes the mean squared error is given by

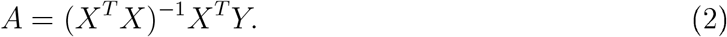

#### Kernelized Ridge Regression (KRR)

Ridge regression is a modification of linear regression. ^34^ Linear regression assumes that there is no relationship between the independent variables. If there is some relationship among the independent variables, ridge regression often gives more reliable results than linear regression.^35^ Ridge regression also has the form *Ŷ*= *XA*, but the matrix *A* is given by

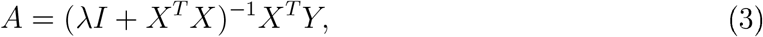

where *λ* ≥ 0 is a complexity penalty and *I* is the identity matrix. The complexity penalty discourages large magnitude entries in *A*, hence reducing overfitting. This equation can also be written as

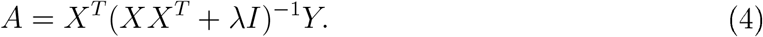

In KRR, the term *XX*^*T*^ is replaced by the Gaussian kernel function, given by

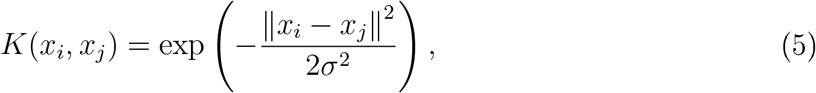

where *σ*^2^ is referred to as the bandwidth of the kernel. The Gaussian kernel measures the similarity between any two points. Given a new input *x*, KRR identifies the portions of the training data with the highest similarity, which therefore have larger influences on the resulting estimate. Let *K* be an *n* by *n* matrix in which the *ij*^th^ entry is *K*(*x*_*i*_,*x*_*j*_), and define

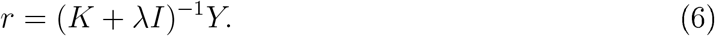

The parameter matrix *A* is given by

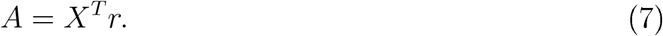

Given new input data *x*, the predicted output is given by

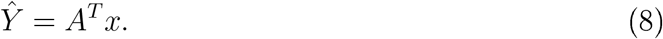

Note that this is equivalently written as

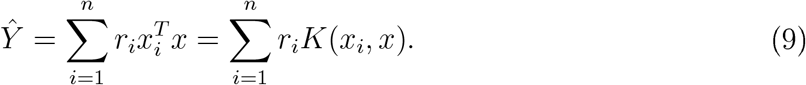

In this study, the Gaussian kernel parameters were found using grid search, as described in Section S2.

#### Decision Tree (DT)

Decision trees recursively partition a given dataset to minimize information entropy. ^34^ Information entropy measures the disorder of a dataset, and is given by

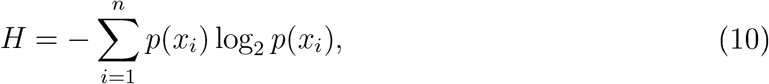

where *p* stands for probability. After each partition, the total entropy of the resulting subsets should be smaller than before. Training a decision tree only requires that one specifies its depth, that is, the level of partition. For example, a tree of depth two splits the original dataset into four subsets. Each final subset is called an end node. A prediction is made by passing a new data point through the decision tree, and outputting the average value of the training data at the resulting end node.

### 2.3. Training and Evaluation Methods

Training is the process of learning model parameters from given (*X, Y*) pairs. In this case, the (*X, Y*) pairs that make up the training data could be obtained in several ways, for example, from traditional counting-based quantification, or from an existing dataset. Evaluation of a model is done by inputting data *X*, then comparing the estimated *Ŷ* with the known *Y* . In this study, the learning algorithms were trained and tested through leave-one out cross-validation. In each dataset of 100 samples, 99 were used for training and the left-out sample was used to test the trained algorithm’s performance. This process was repeated for every sample in each dataset, hence yielding 100 predictions. The accuracy of each algorithm was quantified by mean absolute error (MAE), mean percentage error (MPE) and normalized root mean square error (NRMSE), which are defined as

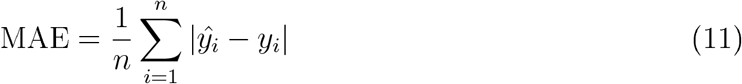

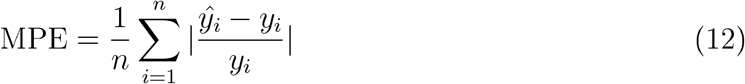

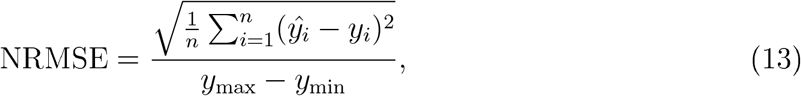

where *y* and *ŷ* are respectively the true and predicted values. There were occasional samples in which the number of a certain type of particle was zero, and these cases were not included in MPE calculation. Another normalization factor, *y*_*max*_−*y*_*min*_, was used to calculate NRMSE to avoid zero denominators. We conducted our numerical experiments as described above, and also with measurement error, which reflects non-idealities such as balance roundoff. We simulated these errors by adding random numbers uniformly distributed between −30% and 30% of the original value to the two finer mesh sieve weights. Random numbers uniformly distributed between −10% and 10% were added to the two coarser mesh sieve weights.

The 95% confidence intervals of the synthetic datasets were calculated using 1000 boot-strapped samples, in order to achieve a normal distribution. The statistical difference between two sets of predictions were evaluated using the Wilcoxon Signed-Rank Test. A significance level of *α* = 0.05 was used.

To test the effect of training dataset size on prediction accuracy, the following number of training samples were also tested: 90, 80, 70, 60, 50, 40, 30, 20, 10, 8, 6, 4, and 2. The corresponding testing dataset sizes were 10, 20, 30 and so on. For each synthetic sample composition, the training-and-testing procedure was repeated 20 times, and each time the training samples were randomly selected. The final result was reported as MPE for each type of particle against the number of training samples.

## 3. Results and Discussion

### 3.1. Training with Synthetic Data

We test the algorithms on the synthetic data described in the Methods Section. The performances of the three algorithms were evaluated by comparing the MAE with 95% confidence interval for the synthetic samples. The results are reported in Table 1. The KRR prediction results were statistically different from LR and DT (p-values *<* 10^*−*8^) The results of LR and DT were similar (p-value = 0.57) Among the three algorithms, KRR always resulted in the lowest average MAE, both with and without incorporating weight measurement error After adding weight measurement error, the performance of each algorithm did not significantly decrease (p-values were 0.71, 0.13 and 0.97 for KRR, LR and DT respectively). This suggests that all algorithms are somewhat robust to weight measurement error. We remark that DT is often poor at predicting continuous values because its output comes from a discrete set of values, the size of which is the number of end nodes.

**Table 1:** Mean Absolute Prediction Error

|  | KRR<sup>a</sup> | LR<sup>b</sup> | DT<sup>c</sup> | KRR-E<sup>d</sup> | LR-E<sup>d</sup> | DT-E<sup>d</sup> |
| --- | --- | --- | --- | --- | --- | --- |
| Rubber | $1.98 \pm 0.07$ | $2.14 \pm 0.07$ | $2.37 \pm 0.10$ | $2.02 \pm 0.08$ | $2.14 \pm 0.07$ | $2.34 \pm 0.09$ |
| Fragment | $2.06 \pm 0.08$ | $2.10 \pm 0.07$ | $2.99 \pm 0.10$ | $2.19 \pm 0.08$ | $2.23 \pm 0.08$ | $3.04 \pm 0.10$ |
| Bead | $3.25 \pm 0.18$ | $3.41 \pm 0.18$ | $4.17 \pm 0.20$ | $3.40 \pm 0.18$ | $3.60 \pm 0.18$ | $4.14 \pm 0.20$ |
| Film | $5.62 \pm 0.25$ | $5.79 \pm 0.23$ | $6.21 \pm 0.25$ | $5.72 \pm 0.23$ | $5.88 \pm 0.25$ | $6.50 \pm 0.28$ |
| Fiber | $20.2 \pm 0.53$ | $20.7 \pm 0.53$ | $22.3 \pm 0.59$ | $20.7 \pm 0.49$ | $21.1 \pm 0.53$ | $22.7 \pm 0.62$ |
| Non-Fiber<sup>e</sup> | $3.67 \pm 0.12$ | $3.74 \pm 0.14$ | $5.07 \pm 0.18$ | $3.89 \pm 0.14$ | $3.97 \pm 0.14$ | $5.10 \pm 0.17$ |

<sup>a</sup> Kernel Ridge Regression; <sup>b</sup> Linear Regression; <sup>c</sup> Decision Tree, implemented with Scikit-Learn<sup>36</sup>; <sup>d</sup> With weight measurement error; <sup>e</sup> Total number of microplastic particles excluding fibers

KRR can be expected to outperform LR because, whereas LR assumes a linear relationship between the data, KRR can accommodate linear and some non-linear relationships. While highly scalable, KRR is more complicated than LR in that one must tune its model parameters (*λ* and *σ*). When simplicity is more important than accuracy, LR may be a superior choice.

We hereon focus on KRR because it achieved the best performance. Its error for the synthetic datasets is summarized in Figure 1. On rubber, KRR achieves low MAE, high MPE and high NRMSE because rubber is usually present in smaller numbers than other particle types. As a result, even an absolute error of one or two particles can result in a percentage error that is more than 50%. On the contrary, fibers are the most numerous. This leads to a large MAE, which, when divided by the large number of fiber particles, leads to a small MPE. For other particles, despite the fact that they are present in similar numbers, films have larger median and average errors than beads, and beads have larger errors than fragments. MAE differs for each size fraction because each size fraction has different amounts of particles. The 500 − 1000 *μ*m size range has the most particles and thus the largest MAE. NRMSE for rubber, fragment, film and fiber is almost constant for all size ranges. Altogether, these results show that the larger the number of microplastics in a given category, the higher its MAE and the lower its MPE.

**Figure 1:**
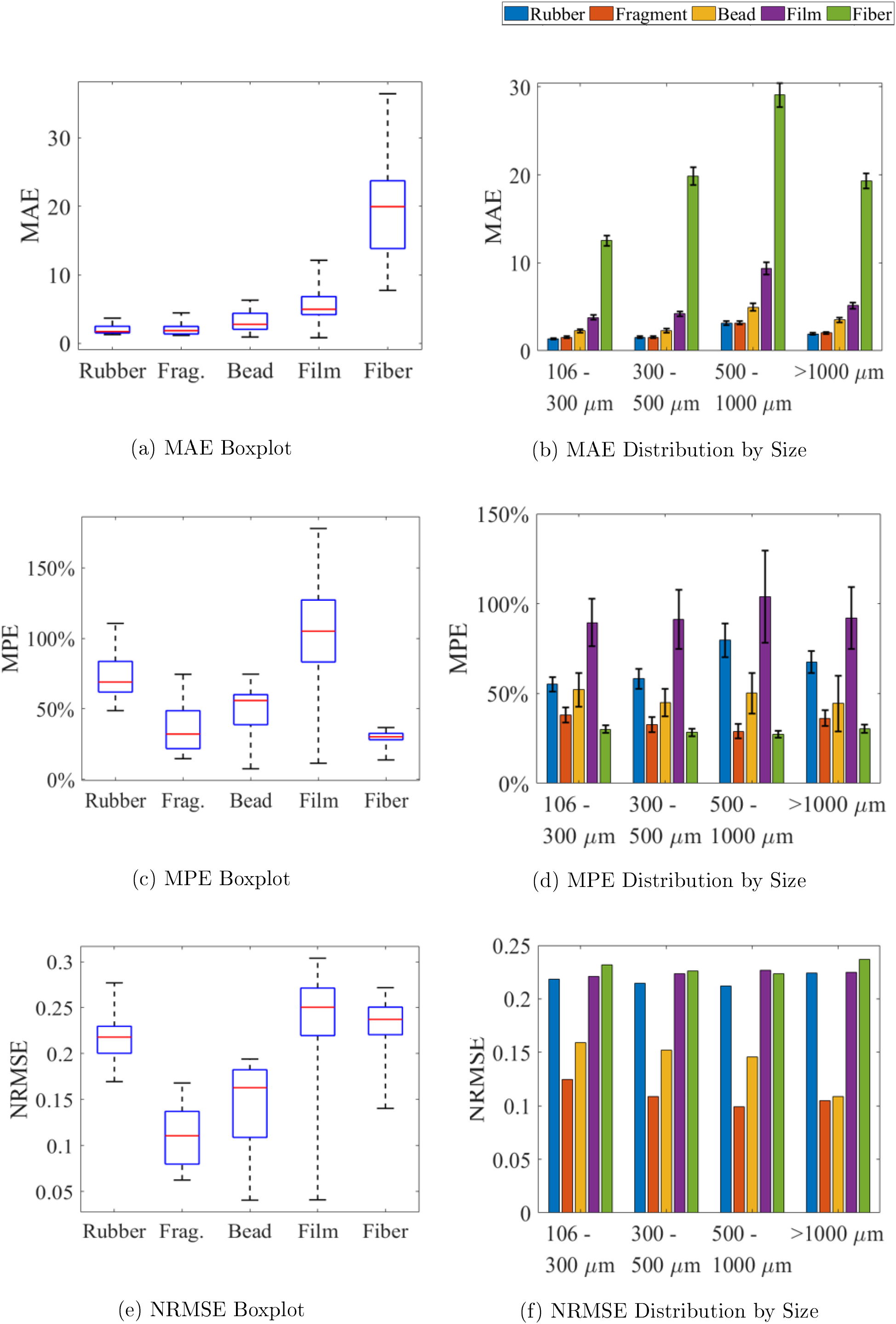
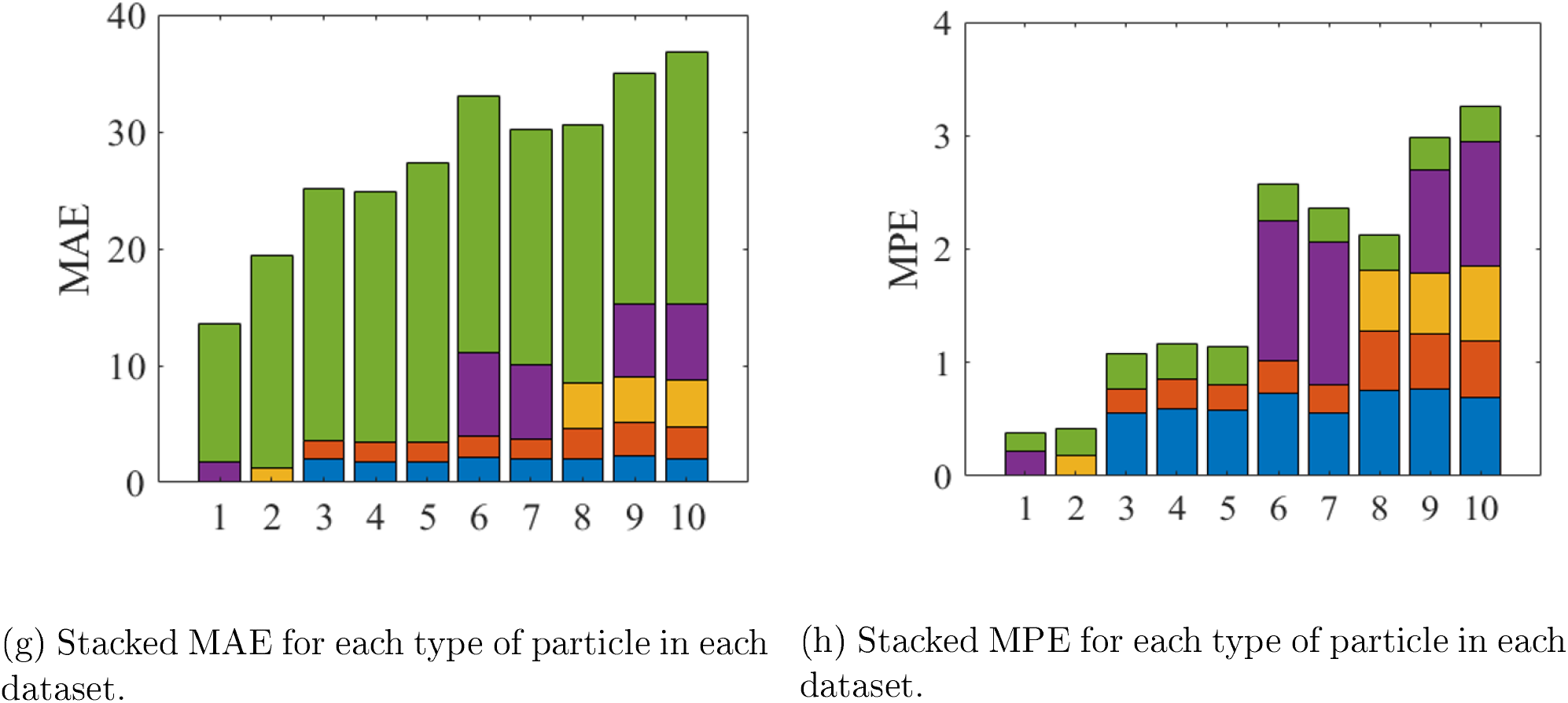
Error of KRR(Kernel Ridge Regression) on synthetic data. In the boxplots, the upper and lower bars represent the upper and lower bounds of the data. The box represents the 25 and 75 percentile, and the middle line represents the median value. Error bars on the right hand side are 95% bootstrapped confidence interval. Frag. represents fragment. Sample complexity increases from composition 1 to 10. MAE: mean absolute error. MPE: mean percentage error. NRMSE: normalized root mean square error.

Figure 1g shows the stacked MAE for each of the ten synthetic datasets. The smallest prediction errors (MAE, MPE and NRMSE) for films and beads occur in their respectively purest samples, e.g., dataset 1 is composed entirely of films and fibers, and dataset 2 of beads and fibers. These values are even lower than that of fragments, because fragments always appear with rubbers in the synthetic datasets. MAE for beads and films increases dramatically with the addition of fragments and rubbers, while MAE for fragments only increases notably with the addition of beads. Also note that the smallest MAE for fibers occurs in dataset 1, made of only films and fibers. This is most likely because films are very light, so that fibers have a higher weight percentage in this dataset than in others. For mixture samples containing fragments, beads and films, MPEs for each type of particles increase with their decreasing weight percentage, as shown in Table S3: average MPE can increase from 14% to 51% for fragments if their weight percentage drops from 80% to 20%. All these observations suggest that, when aggregate weight is used as the sole feature, heavier particle types have consistently lower prediction error because of their higher weight percentage in each sample. We can also see that MAE does not change significantly after adding various amounts of organics, most likely because the total weight of organics is smaller than that of the plastics, and the addition of organics does not change the order of weight percentages of other particles. For samples severely contaminated by organic particles, e.g., a lake with large amount of algae or plant debris sampled along with microplastics, an organic digestion step will be necessary prior to drying and measuring the sample weight, in order to limit the contribution of organics to the total sample weight.

To summarize, the procedure is accurate for particles that account for a significant fraction of a sample’s weight. This is why it is relatively difficult to predict fiber counts in mixed samples, which are numerous but might account for only a few percent of the overall weight. On the other hand, it is relatively easy to predict fragment counts in mixed samples because they account for a moderate fraction of the overall weight.

The total error associated with microplastic count prediction by KRR remains comparable to or lower than that of manual counting, even for the more heterogenous samples. Isobe *et al*.^23^ used artificial samples to estimate the relative uncertainty of microplastic counts performed by humans. In their study, samples with *n*_0_ ≈ 90 fragments less than 2 mm were sent to 12 different laboratories. The relative error of a given laboratory s count, *n*, was calculated as (*n* − *n*_0_)*/n*_0_. The average error of all laboratories was around ±50%, which we interpret as the error of a typical human count. In other visual sorting microplastic studies, researchers have also reported a false positive ratio ranging from 20% to 70%, meaning that, among the particles visually sorted as microplastics, 20% to 70% of them were determined via spectroscopic analysis to be non-plastic. ^8,29,37,38^

In our synthetic datasets, the average number of fragments, beads and films is around 50 for each, whenever present. Therefore, based on the similarity of *Interlab* samples to our synthetic data, it is reasonable to compare our results with theirs, with respect to each particle type. As discussed earlier, when these particles are present in their respective purest samples, their MPE can be less than 10%. In mixed samples, however, MPE for beads can go beyond 60% and films can go beyond 100%. Figure 1h shows MPE for each type of particle in the ten synthetic datasets. In mixed samples, MPE for films is always above 90%, due to their low contribution to the total weight, and that for rubbers is always above 50%, due to their small amount in each sample. MPE for fragments and fibers is below 35% except for datasets 8, 9, and 10, where the presence of beads increases the prediction error for fragments. When all particles are present, e.g., datasets 9 and 10, the prediction accuracy is better for fibers, comparable for beads and fragments, and worse for films and rubbers, compared to the accuracy of visual sorting. However, MPE decreases with increasing sample size, as indicated in Table S4. As the number of each type of particle increases, MPE for fragment, bead and film gradually decreases. MPE for fibers does not change significantly, but MPE for rubber increases notably. This is because the number of rubber particles is a fraction of the number of fragments, as stated in Table S2. Then, there are several cases where the number of rubber particles is small, and the prediction is large, resulting in a large average MPE.

The above leave-one-out experiments correspond to training data with 99 samples, which may be impractically large in some cases. The effect of the number of training samples on KRR prediction accuracy can be found in Section S4. In general, MPE bottoms out when the number of training samples exceeds 20, regardless of sample composition and particle type. Therefore, in this case, there is little further accuracy to be gained from having more than 20 samples in the training data, and the leave-one-out results can be seen as the typical KRR performance.

These findings suggest that KRR, if given a moderate amount of accurate training data, can estimate counts more accurately than visual sorting on samples with homogeneous composition, on heavier particles in mixed samples, and on larger samples, i.e., samples with larger particle number and total weight.

### 3.2. Training with Experimental Data

Figure 2 shows the performance of KRR on five datasets obtained from experiments: *Interlab, Washing Machine, Cocos, Shearwaters*, and *Great Lakes*. Training and testing was done through leave-one-out cross-validation. Data preprocessing was performed only for the *Great Lakes* dataset. For the latter, we first merged the 11 types of morphologies from the original report down to 6 (fiber, film, foam, fragment, pellet and rubber) in order to reduce data sparsity. Then, 3 outliers (outside three times the interquartile range of all data) were excluded from training and testing. Finally, training and testing was performed with the number of each type of particle, and with the total number of all particles, separately. Figures 2a to 2f show the predicted values plotted against the measured values. In most cases, the outcomes scatter closely around the diagonal line, which represents perfect prediction. The MPE for all datasets are mainly below 50%, with median values at 25% for *Washing Maching*, 13% for *Interlab*, 32% for *Cocos* and 0% for *Shearwaters*, as shown by Figure 3a. The MPE for the *Great Lakes* dataset varies for different types of particles (Figure 3b), but in most cases the median value is less than 50%. MPEs for fiber, film and rubber are low in this case because the numbers of these particles show small variances, as demonstrated by Figure S16(b). MPEs for pellet, fragment and foam show a similar trend as found with artificial datasets: prediction error is smaller for heavier particles. These results demonstrate that predictions from a well-trained KRR can be more accurate than the results of traditional counting. However, there are a few large prediction errors, which could be the result of patterns undetected by KRR, or errors in the measured counts.

**Figure 2:**
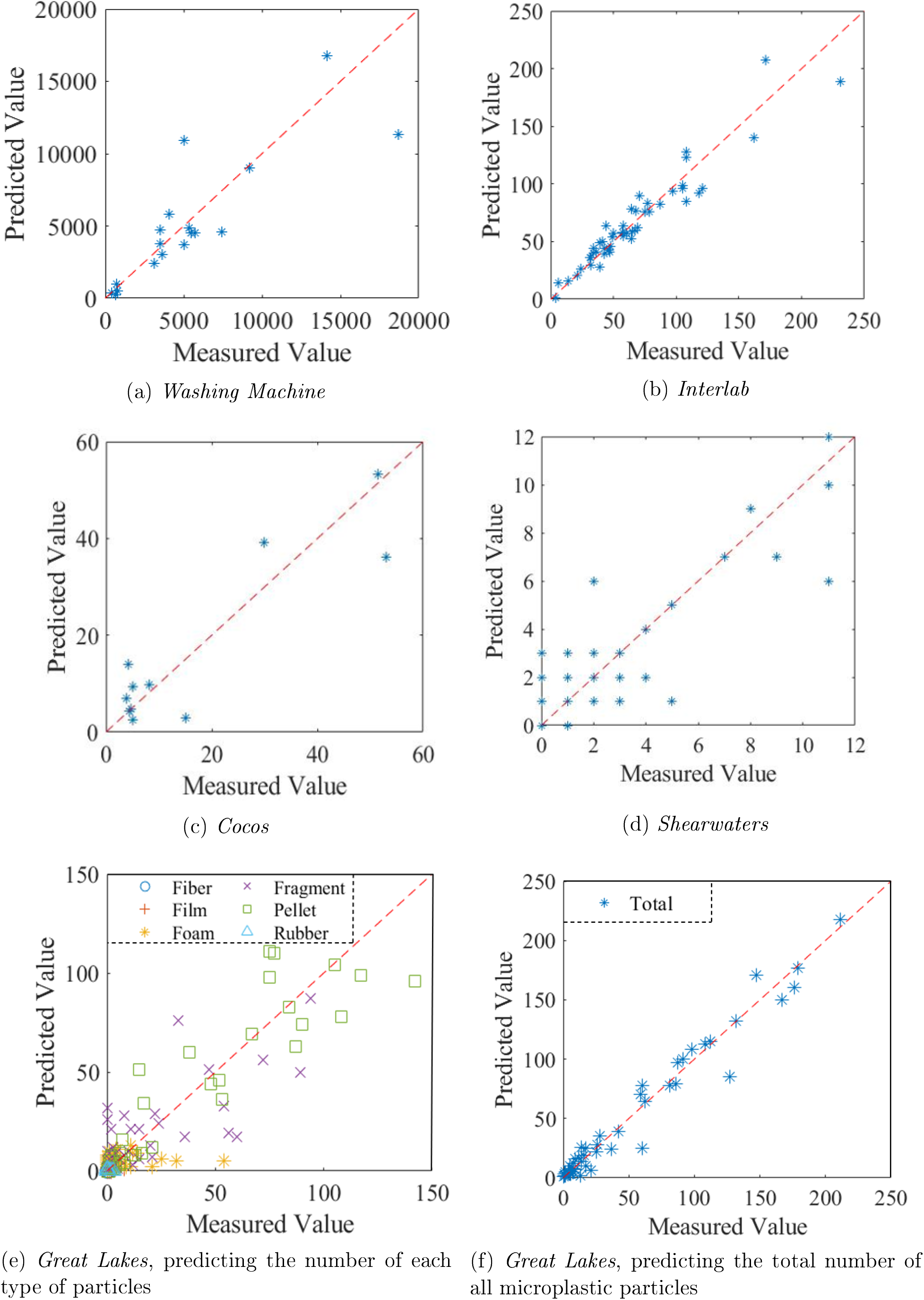
KRR (Kernel Ridge Regression) Performance Summary on Real Data

**Figure 3:**
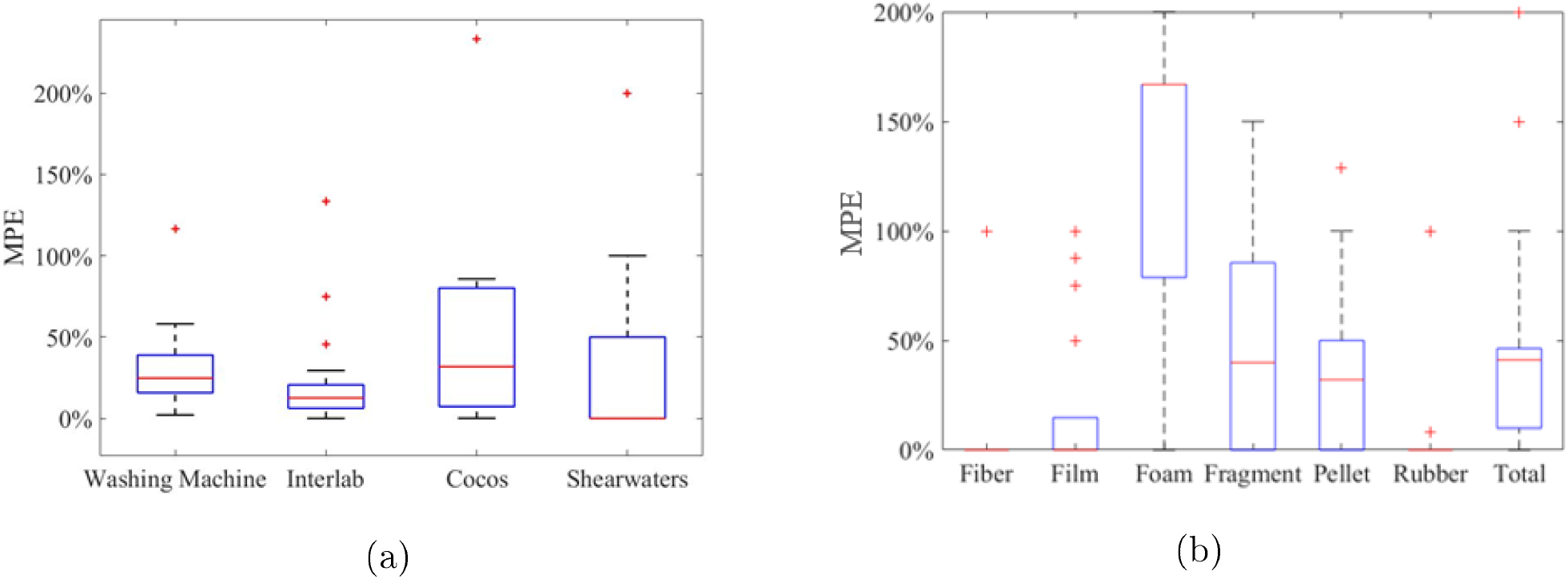
MPE (Mean Percentage Error) of KRR (Kernel Ridge Regression) prediction on: (a) the *Interlab, Washing Machine, Cocos*, and *Shearwaters* datasets, and (b) the *Great Lakes* dataset, with the display of y-axis limited at 200%. Original version is included as Figure S16(a). In the boxplots, the upper and lower bars represent the upper and lower bounds of the data. The box represents the 25 and 75 percentile, and the middle line represents the median value.

If the original count error is Gaussian with zero mean and constant variance, it will not affect the expected value of the model parameters. ^39^ In the case of microplastic quantification, it is possible for human counts to be biased, violating the zero-mean requirement, e.g., by always over-counting cotton fibers from air contamination or ignoring a type of particle thought to be non-plastic. To reduce the effect of biased measurement errors, training datasets should be as accurate as possible, for example, verified by FTIR or Raman spectroscopy and corrected to report plastic-only particles. Most studies that used such methods to confirm particle identities (plastic vs. non-plastic), showed that between 20% to 70%^8,29,37,38^ of the total number of microparticles are not plastics, and for sediment samples this rate can be as high as 98%.^40^ As a consequence, accurate training data is necessary for making accurate predictions.

To optimize prediction quality, a minimum training dataset size is required. The recommended training dataset size increases with the number of predicted variables. When the number of predicted quantities is small, e.g., only fiber counts were predicted for the *Washing Machine* data, a training dataset with 8 or more samples can be adequate, as suggested by Jenkins *et al* .^41^and shown in Figure S15. In cases like synthetic dataset 9, which has 4 size fractions and 5 types of particle, a training dataset of 20 samples is recommended, as demonstrated by our numerical experiments. The training samples should cover as much the data range as possible, meaning that, the samples should be representative of the full range of inputs and outcomes.

### 3.3. Implementation

Assessing the extent and impact of microplastic contamination requires the collection of samples at high temporal and spatial resolutions. This section briefly compares our approach to standard quantification procedures and demonstrates its power to quickly collect large amounts of data. Figure 4 shows the two quantification procedures after organic digestion, density separation, and sieving. This comparison mainly focuses on time, cost and level of expertise required. The equipment needed for traditional manual counting method costs approximately 1, 000 USD for visual sorting. The cost of prediction based on sieved mass is approximately 5, 000 USD to 7, 000 USD in equipment for sample mass determination. Commercial laboratories charge between 1, 000 to 5, 000 USD for microplastic sample analysis, and around 50 to 250 times less for total suspended solid concentration measurement (∼ 20 USD), which is comparable to the proposed method on determining the weight of microplastics. The level of expertise required was estimated based on the fact that visual sorting of microplastic particles requires a certain amount of training; whereas, drying, weighing, and running a trained algorithm are all relatively straightforward tasks. Finally, our method requires between 10% to 40% the time of traditional methods. It can achieve better MPE than manual counting results reported by Isobe *et al*. for heavier particles in a complicated sample, and our MPE for lighter particles can be reduced to below 30% when the sample size increases. ^23^ In conclusion, by using machine learning to predict microplastic counts, we can reduce the experimental labour by 50% to 80%, without significantly increasing the cost or sacrificing reliability.

**Figure 4:**
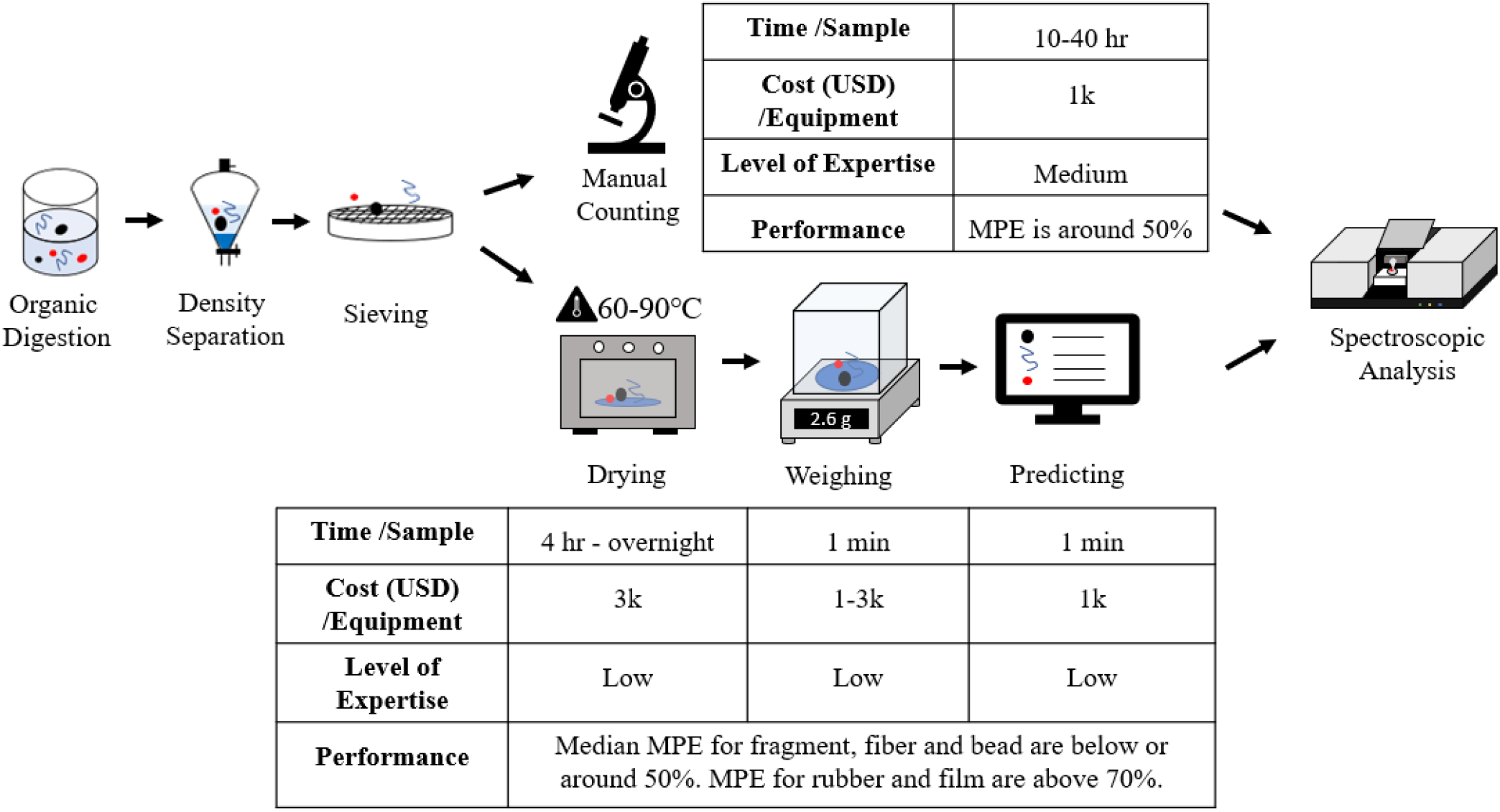
Comparison between the current and the proposed method. k stands for thousand. The time needed to manually count microplastic particles using a microscope was estimated from laboratory experiences, at around 10 to 40 hours per sample, with each sample containing around 50 to 100 particles. Predicting counts from sample weights consists of three parts: 1) drying, which ranges from 4 hours to overnight depending on the temperature used, 2) weighing, estimated as 1 minute, and 3) prediction, estimated as 1 minute for an already trained algorithm. ^24,42^ To this, one can add the time needed to process the required number of samples through the traditional manual counting method to generate a training dataset, i.e., an extra 10 to 40 hours per training sample. The equipment cost for manual counting is dominated by the microscope, estimated as 1,000 USD. The cost for the prediction method is estimated as the total cost of the oven (∼ 3, 000 USD), balance (∼ 1, 000 to 3, 000 USD) and computer (∼ 1, 000 USD). The cost for the balance can be lower for heavier samples where two decimals are enough; whereas, for lighter samples a four-decimal balance will be preferred.

We now discuss three hypothetical use cases. Graphical representations can be found in Section S6. Depending on the context, additional features that might influence the count can be added. When assessing environmental contamination by microplastics, these features can include time of the year, season, longitude, latitude, distance to the nearest urban or industrial center, and so on. We note, however, that a feature will only improve performance if it is actually related to the count. For example, location may be particularly useful if some of sampling sites are near a highway.

In the first hypothetical use case, microplastic contamination is monitored at a single site, e.g., a lake, over the course of a year (Figure S17). If one collects several samples each month, one sample from each collection period can be counted using the traditional method and verified by spectroscopic analysis to form a training dataset. This training data would be used to train KRR. The rest of the samples would be sieved and weighed, and the microplastic counts would be estimated using the trained KRR. Here, the time of the year could be added as an extra feature if it influences microplastic counts.

In the second hypothetical use case, one wishes to monitor a large geographical area (Figure S18). One could perform traditional counting on samples from a small number of locations, use this data to train KRR, and then use the algorithm to quickly estimate counts at a larger number of remaining locations. In this case, location might be included as an additional feature in the training data.

The third hypothetical use case involves samples with homogeneous composition, e.g., atmospheric or washing machine effluent samples composed of mainly fibers (Table S5). As demonstrated before, KRR would perform better if the sample contains only one type of particle. Ideally, one would weigh all collected samples and choose from them several samples that depict the range of the measurements, count them using traditional manual method and use them to train KRR. The rest of the samples can be predicted using the trained model.

## 4. Conclusion

This study demonstrated the possibility of using regression methods for microplastic quantification based on sieved weight measurements. The method works better for pure samples made of one type of particles, or for heavier particles in mixed samples. Note that these results are based on a set of mass measurements, and that further improvement may be attainable by incorporating features such as geographic location and time of measurement. This method is faster and easier than current visual quantification methods. As shown in Figure 4, the new method takes less than half of the time needed for manual counting, and it only involves fundamental laboratory equipment. We therefore believe that it could be widely useful to both experts and non-experts. The accuracy of predictions from KRR are better than that attained by human counting, suggesting that this approach has the potential to accelerate experimental research on this topic without sacrificing the quality of microplastic quantification.

## Acknowledgenents and Data Availability

This research was supported by the University of Toronto and the Natural Sciences and Engineering Research Council of Canada, Canada Research Chairs, grant no. 950-230892. Raw data were generated at the University of Toronto. Derived data supporting the findings of this study are available from the corresponding author E.P. on request.

## Graphical TOC Entry

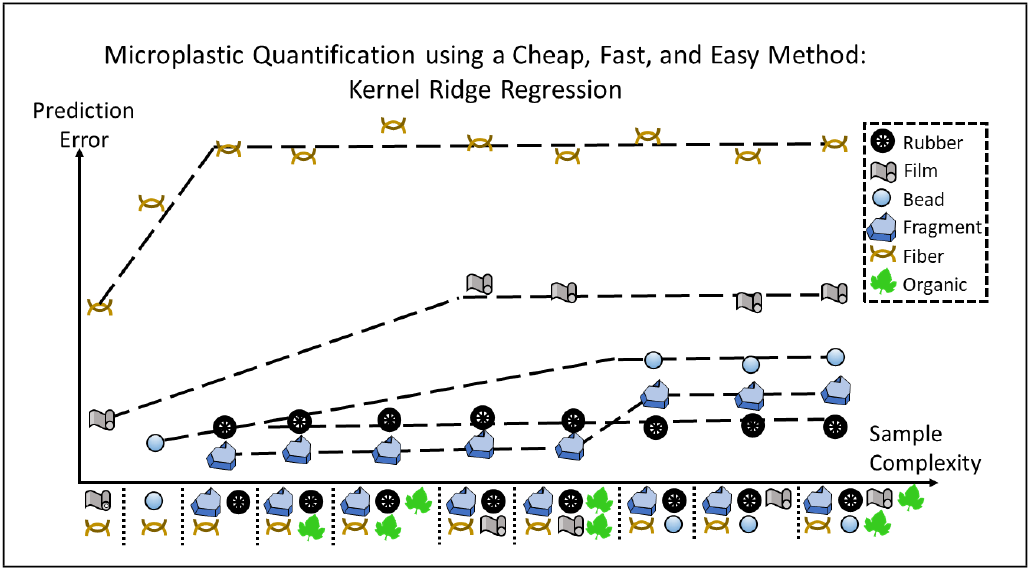

## Highlights

- Predict microplastic counts based on aggregate mass measurements
- Better prediction accuracy for homogeneous samples and heavier particles
- Mass measurement error did not affect prediction accuracy
- Faster, cheaper and easier microplastic quantification method

## Notes

### Competing Interest Statement

The authors have declared no competing interest.

