## Supporting Information for "Efficient Prediction of Microplastic Counts from Mass Measurements"

Shuyao Tan[1], Joshua Taylor[2], and Elodie Passseport[*1,3]

[1]Department of Chemical Engineering and Applied Chemistry, University of Toronto

[2]The Edward S. Rogers Sr. Department of Electrical & Computer Engineering, University of Toronto

[3]Department of Civil and Mineral Engineering, University of Toronto

### Section S1: Information about the Synthetic Datasets

**Table S1: Synthetic Data Generation Parameters**

| Particle Type | Shape | Number Range | Density Range (g/cm$^3$) |
|---|---|---|---|
| Rubber | Cuboid | 0-50% of F[a] | 0.4-0.6[1] |
| Fragment | Cuboid | 0-100 | 0.8-1.4[2] |
| Film | Cuboid | 0-100 | 0.8-1.4[2] |
| Bead | Sphere | 0-100 | 0.8-1.4[2] |
| Fiber | Cylinder | 100-600 | 1.1-1.4[3] |
| Organics | Cuboid | Various | 0.1-1.3[4] |

[a] F stands for the number of fragments. The total number of fragment and rubber particles was between 0 and 100. Between 0 and 50% of them were rubber particles, and the rest were fragments.

Reference for Table S1:

(1) *User guidelines for waste and byproduct materials in pavement construction* ; U.S. Department of Transportation, 2016.

(2) *Microbeads - A Science Summary*; Environment and Climate Change Canada, 2015.

(3) Kooi, M.; Koelmans, A. A. Simplifying Microplastic via Continuous Probability Distributions for Size, Shape, and Density. *Environmental Science & Technology Letters* **2019**, *6*, 551-557.

(4) Niinemets, Ü. Research review. Components of leaf dry mass per area - thickness and density - alter leaf photosynthetic capacity in reverse directions in woody plants. *New Phytologist* **1999**, *144*, 35-47.

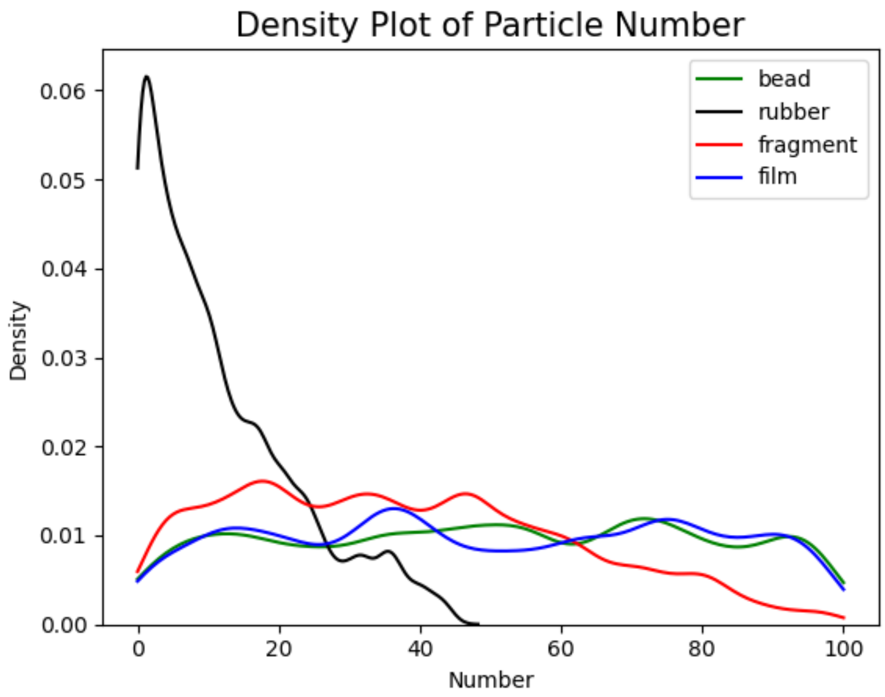


**Figure S1: Probability Densities of Bead, Rubber, Fragment and Film**

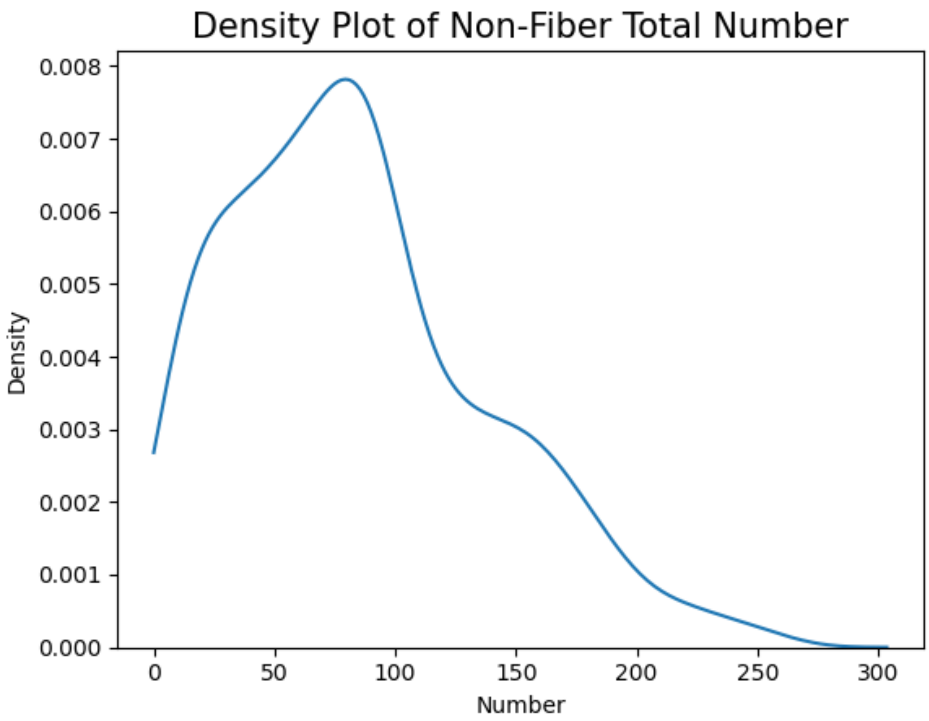


**Figure S2: Probability Density of Total, Non-Fibrous Particles**

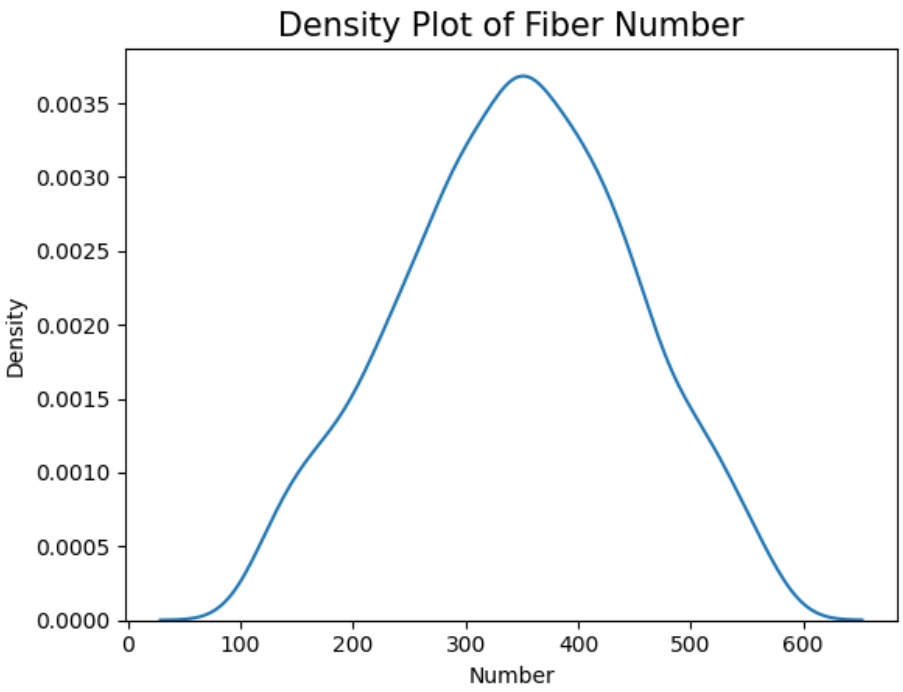


**Figure S3: Probability Density of Total Fibers**

**Table S2: Synthetic Data with Larger Samples**

| Particle Type | Twice Particles | Four Times Particles |
|---|---|---|
| Rubber | 0-50% of F[a] | 0-50% of F[a] |
| Fragment | 50-200 | 200-400 |
| Film | 50-200 | 200-400 |
| Bead | 50-200 | 200-400 |
| Fiber | 200-1200 | 400-2400 |

[a] F stands for the number of fragments. The total number of fragment and rubber particles was between 0 and 100. Between 0 and 50% of them were rubber particles, and the rest were fragments.

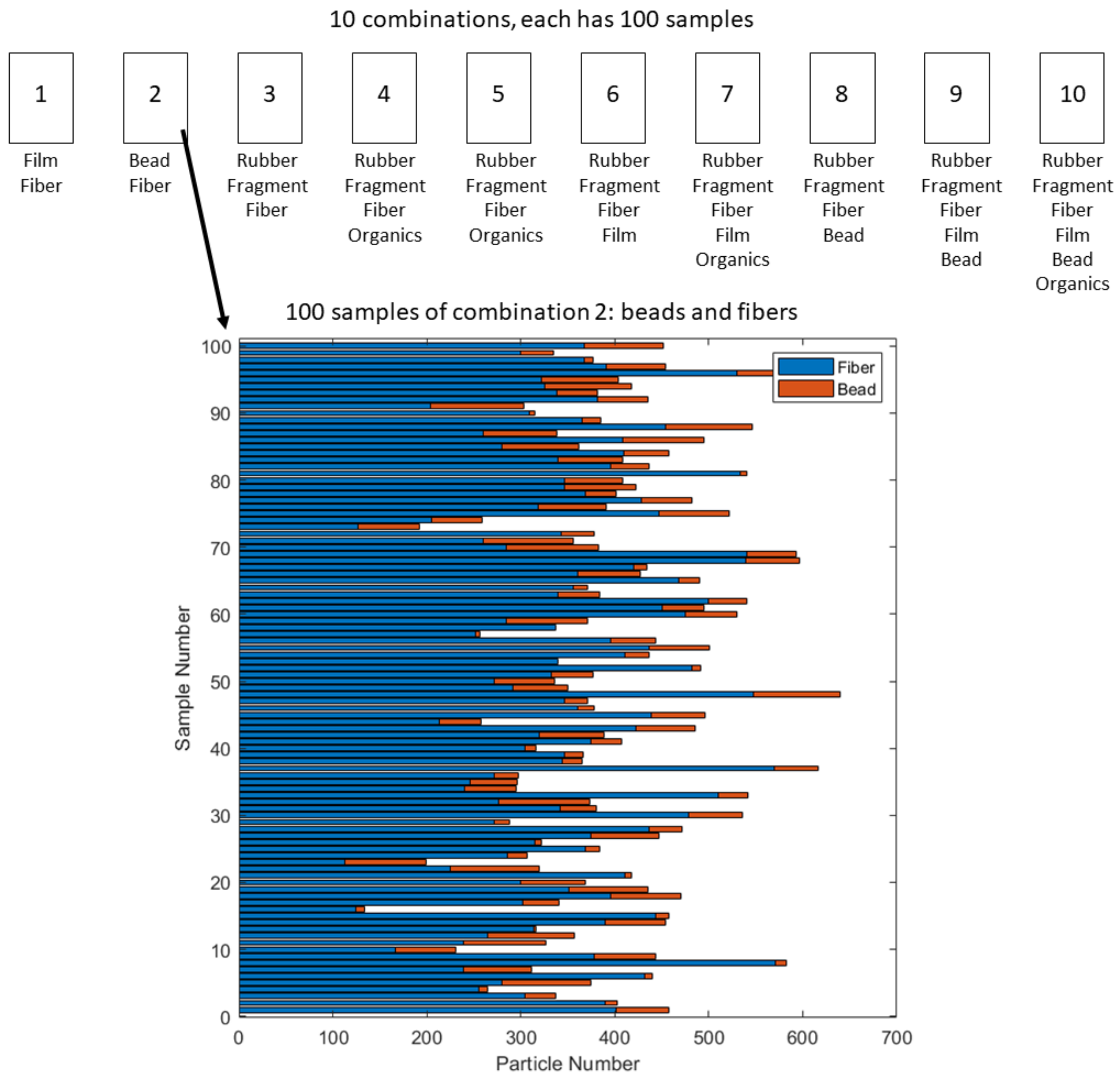


**Figure S4: Example of 100 Synthetic Samples**

**Section S2: Gaussian Kernel Parameters**

The penalty factor is chosen from:

[1.e-11, 1.e-10, 1.e-09, 1.e-08, 1.e-07, 1.e-06, 1.e-05, 1.e-04, 1.e-03, 1.e-02, 1.e-01, 1.e+00, 1.e+01, 1.e+02, 1.e+03]

The bandwidth is chosen from:

[1 1.78 3.16 5.62, 10, 17.78, 31.62, 56.23, 100, 177.83, 316.23, 562.34, 1000, 1778.3, 3162.3]

**Section S3: Performance of KRR on Various Samples**

**Table S3: Effect of particle weight percentage on MPE**

| Weight Percentage (%) | <20 | 20-50 | 50-80 | >80 |
|---|---|---|---|---|
| Fragment MPE (%) | 51.4 | 31.9 | 17.7 | 14.1 |
| Bead MPE (%) | 57.3 | 40.9 | 25.9 | 18.2 |

**Table S4: Kernel ridge regression mean percentage error with increased particle number, all numbers shown are percentages**

| Particle Type | Base Case | Twice Particles | Four Times Particles |
|---|---|---|---|
| Rubber | 76 ± 12 | 102 ± 17 | 137 ± 37 |
| Fragment | 49 ± 6.9 | 30 ± 3.7 | 18 ± 1.8 |
| Film | 91 ± 16 | 42 ± 4.5 | 20 ± 1.8 |
| Bead | 54 ± 13 | 29 ± 4.7 | 15 ± 1.3 |
| Fiber | 28 ± 3.2 | 33 ± 3.5 | 26 ± 2.9 |

**Section S4: Effect of Training Dataset Size**

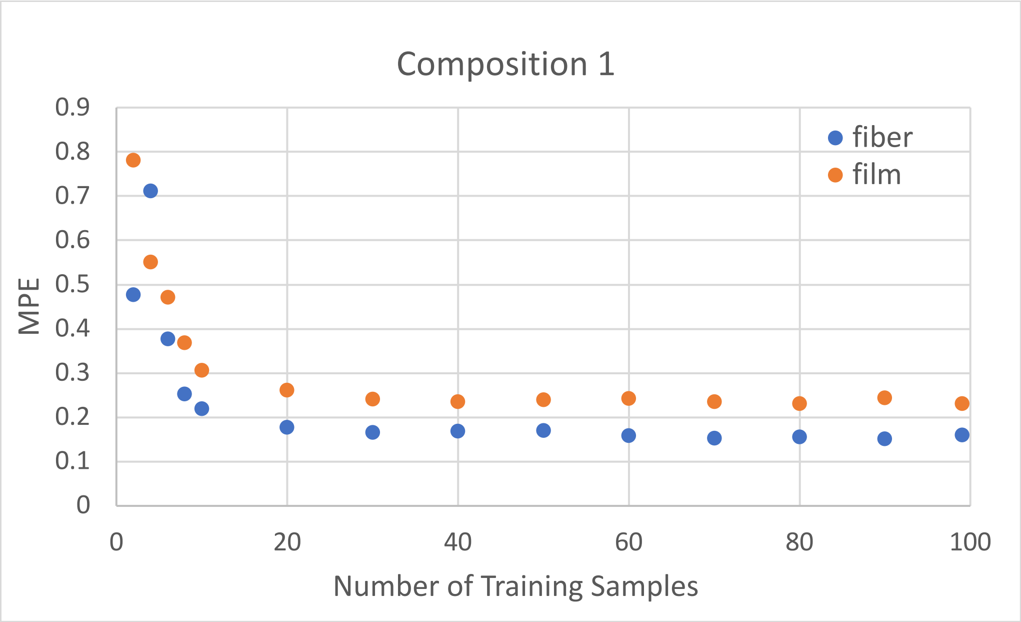


**Figure S5(a): Effect of Training Dataset Size on KRR Prediction Accuracy, Composition 1, Average Value**

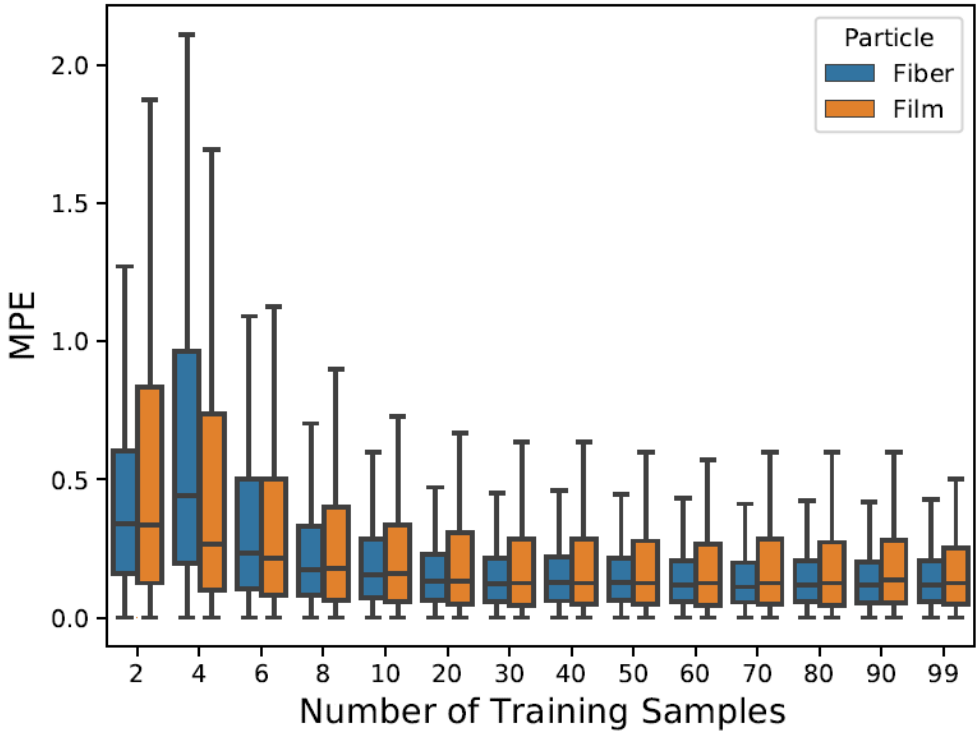


**Figure S5(b): Effect of Training Dataset Size on KRR Prediction Accuracy, Composition 1, Boxplot**

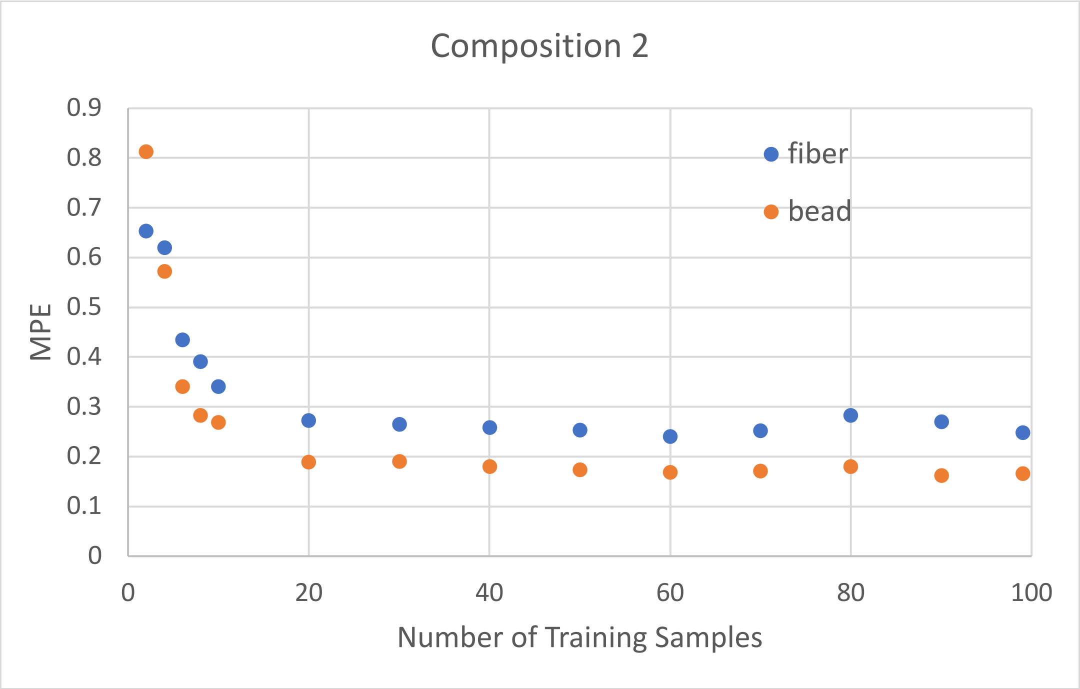


**Figure S6(a): Effect of Training Dataset Size on KRR Prediction Accuracy, Composition 2, Average**

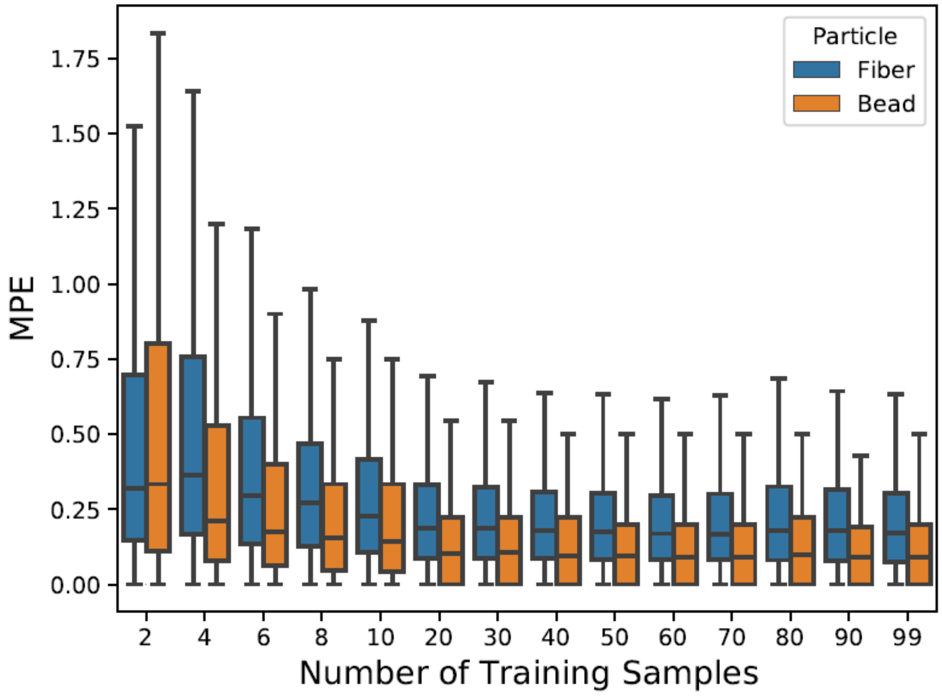


**Figure S6(b): Effect of Training Dataset Size on KRR Prediction Accuracy, Composition 2, Boxplot**

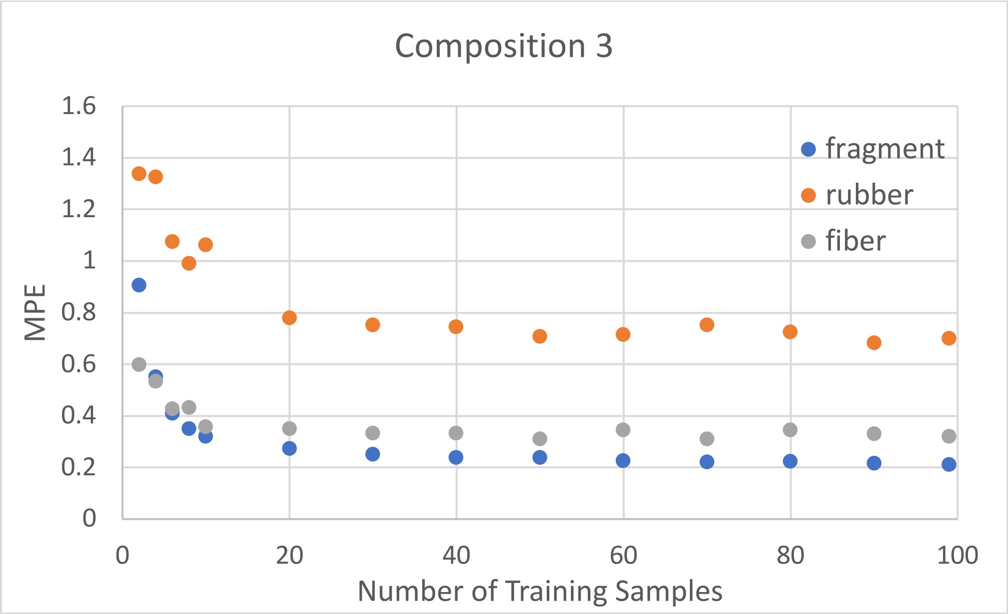


**Figure S7(a): Effect of Training Dataset Size on KRR Prediction Accuracy, Composition 3, Average**

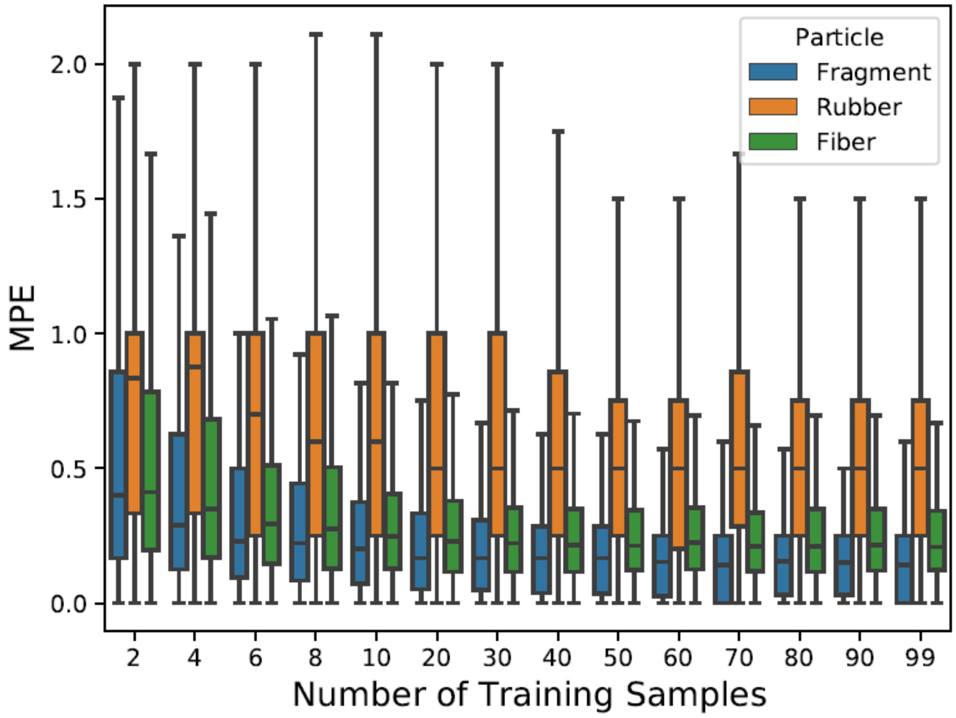


**Figure S7(b): Effect of Training Dataset Size on KRR Prediction Accuracy, Composition 3, Boxplot**

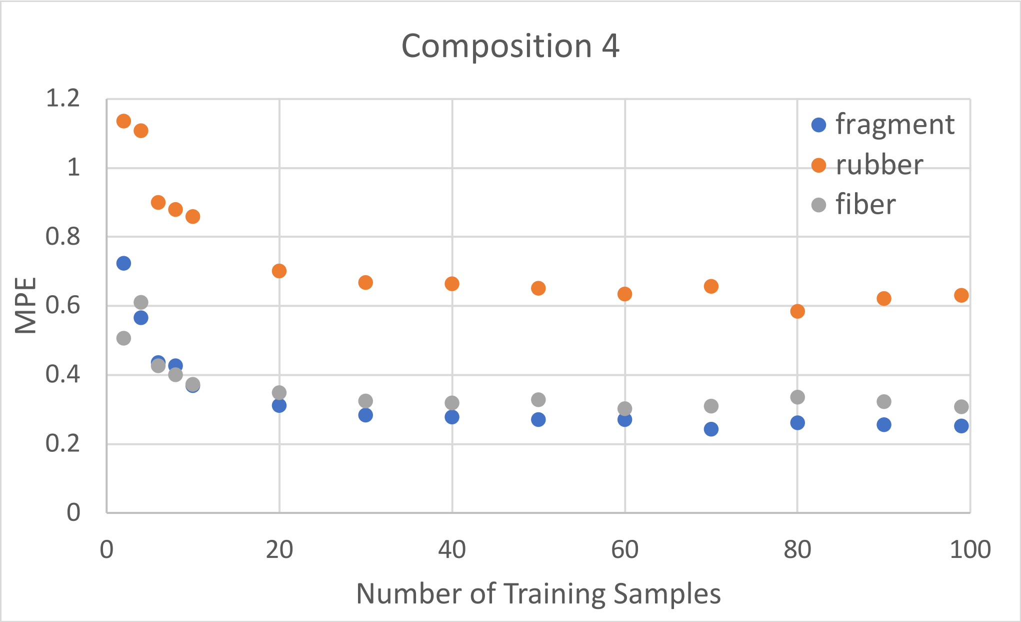


**Figure S8(a): Effect of Training Dataset Size on KRR Prediction Accuracy, Composition 4, Average**

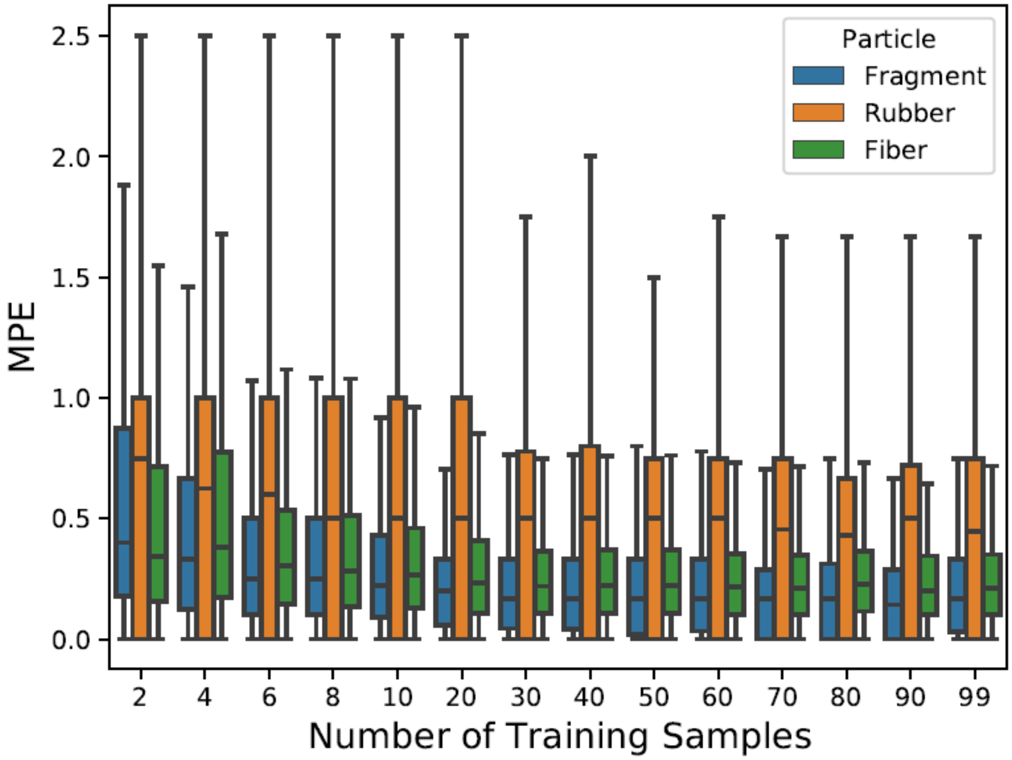


**Figure S8(b): Effect of Training Dataset Size on KRR Prediction Accuracy, Composition 4, Boxplot**

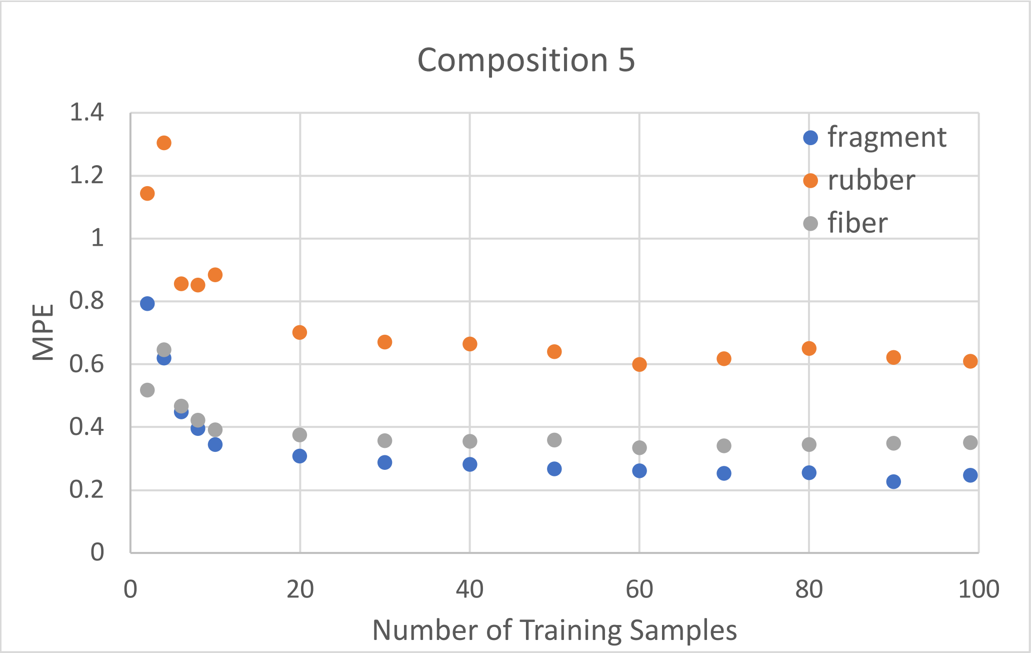


**Figure S9(a): Effect of Training Dataset Size on KRR Prediction Accuracy, Composition 5, Average**

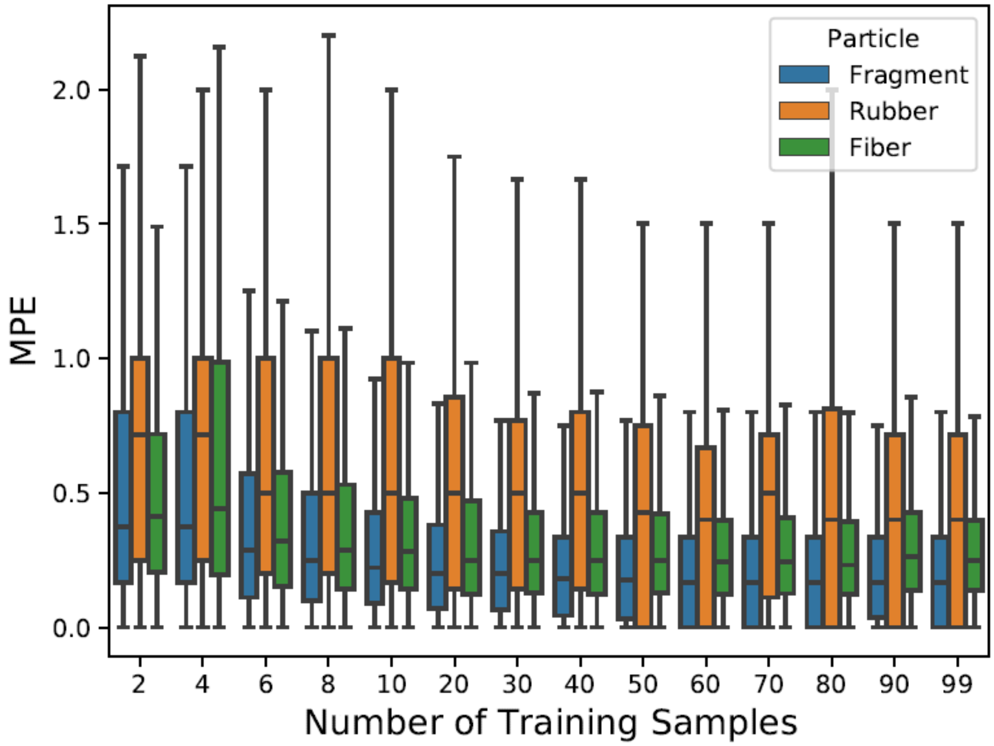


**Figure S9(b): Effect of Training Dataset Size on KRR Prediction Accuracy, Composition 5, Boxplot**

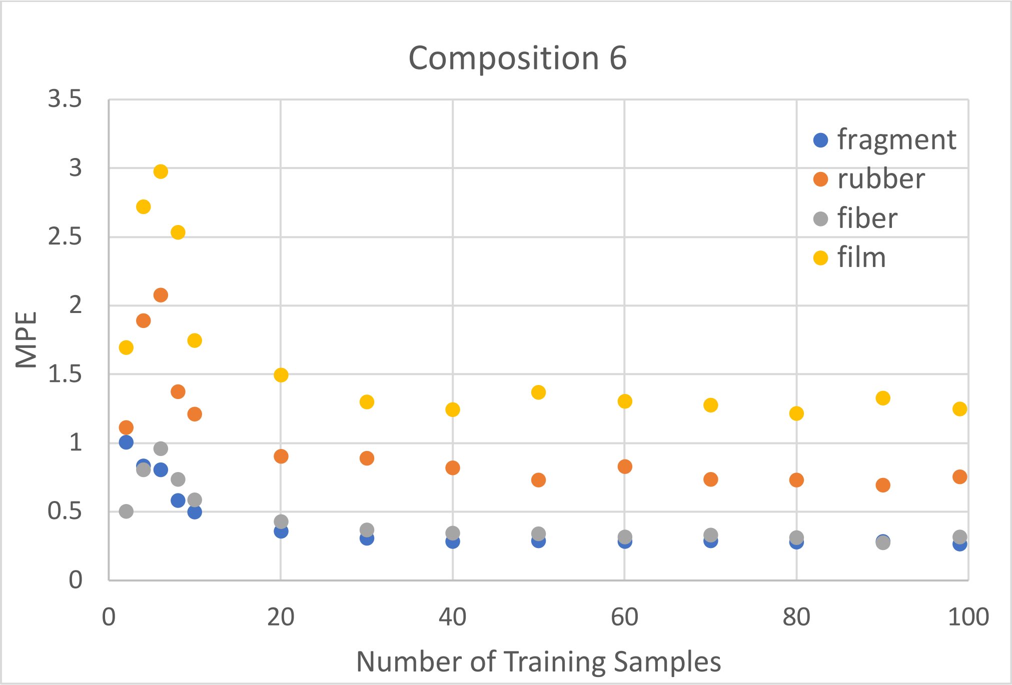


**Figure S10(a) : Effect of Training Dataset Size on KRR Prediction Accuracy, Composition 6, Average**

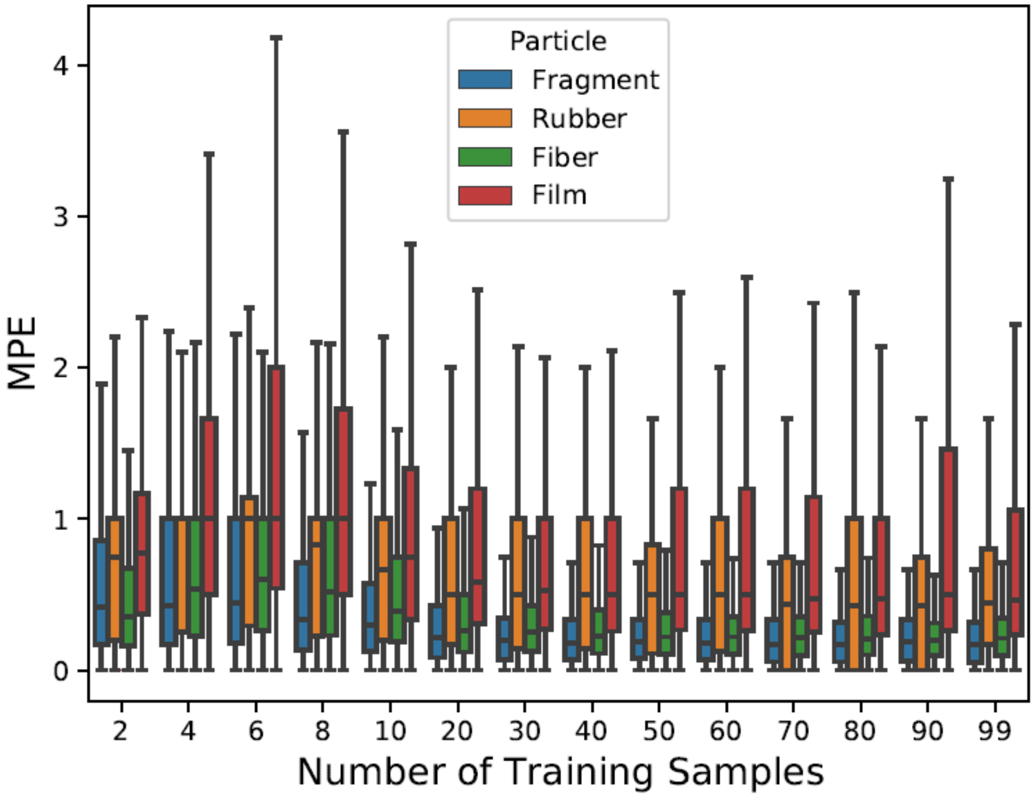


**Figure S10(b) : Effect of Training Dataset Size on KRR Prediction Accuracy, Composition 6, Boxplot**

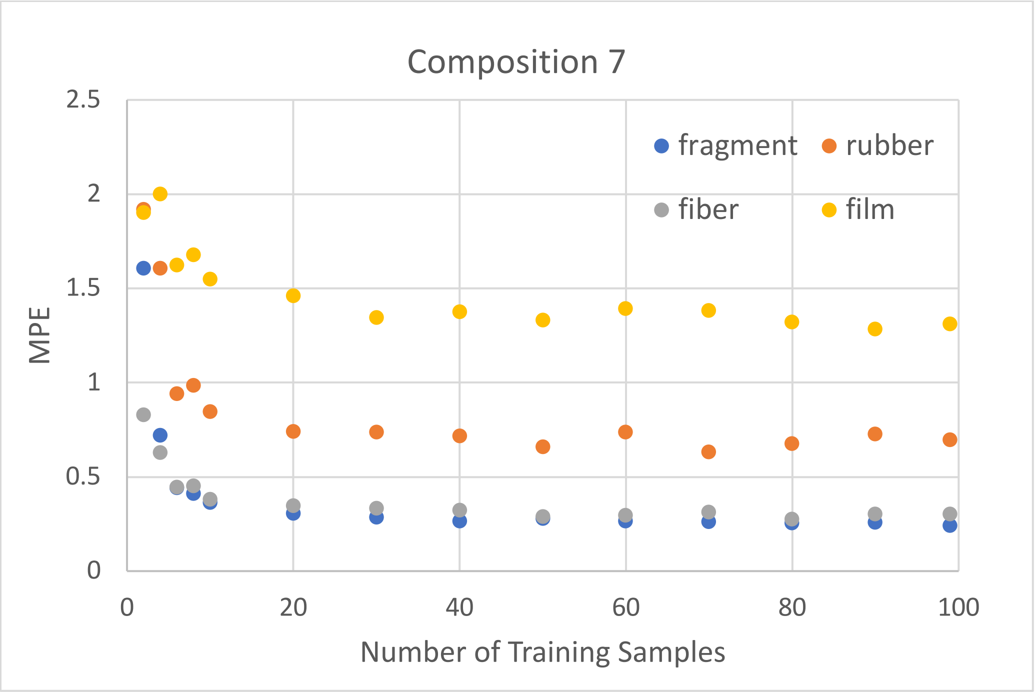


**Figure S11(a): Effect of Training Dataset Size on KRR Prediction Accuracy, Composition 7, Average**

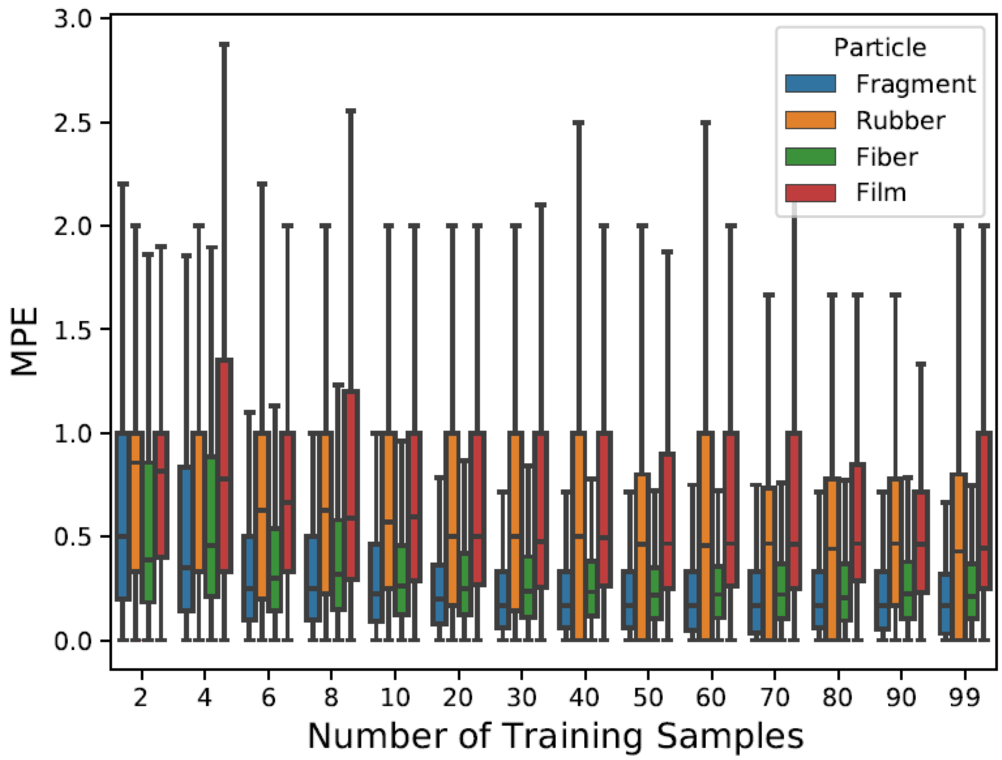


**Figure S11(b): Effect of Training Dataset Size on KRR Prediction Accuracy, Composition 7, Boxplot**

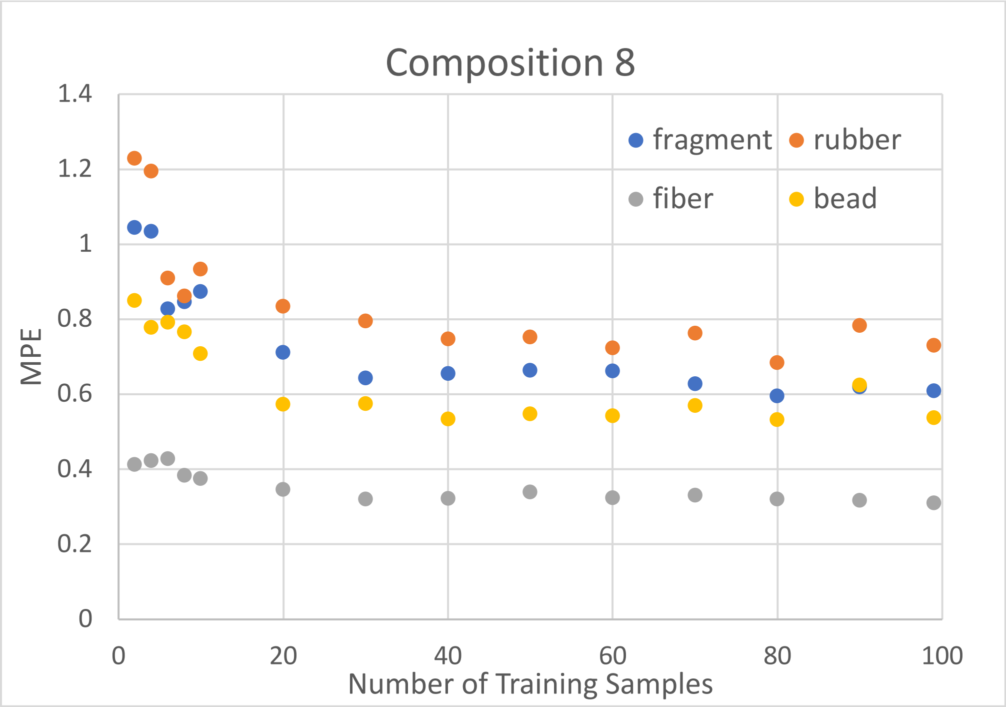


**Figure S12(a): Effect of Training Dataset Size on KRR Prediction Accuracy, Composition 8, Average**

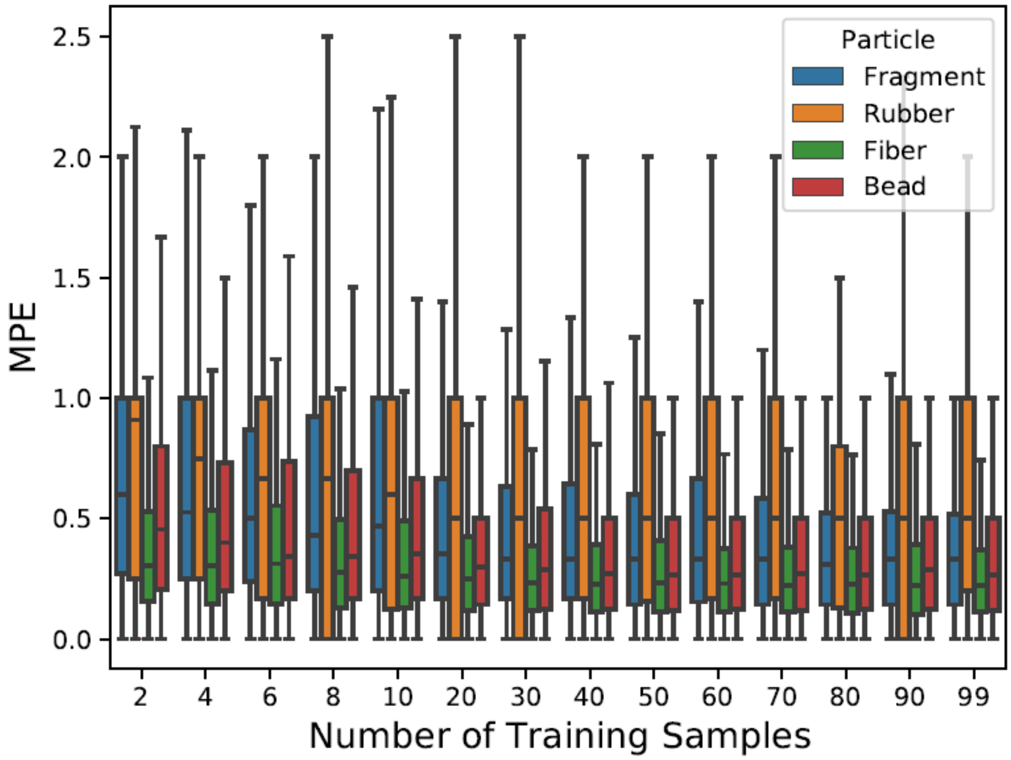


**Figure S12(b): Effect of Training Dataset Size on KRR Prediction Accuracy, Composition 8, Boxplot**

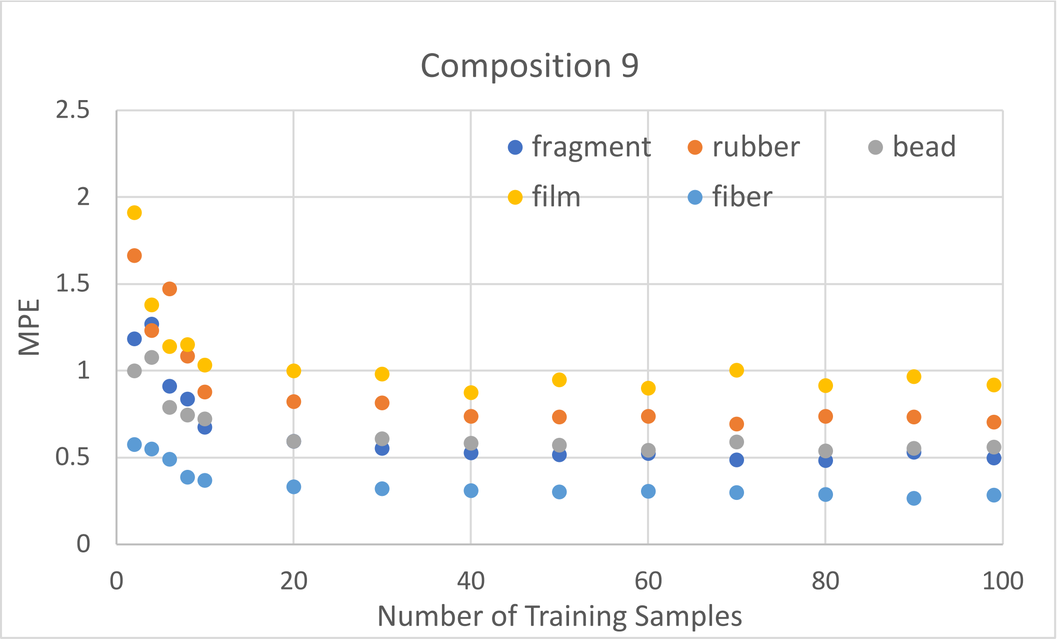


**Figure S13(a): Effect of Training Dataset Size on KRR Prediction Accuracy, Composition 9, Average**

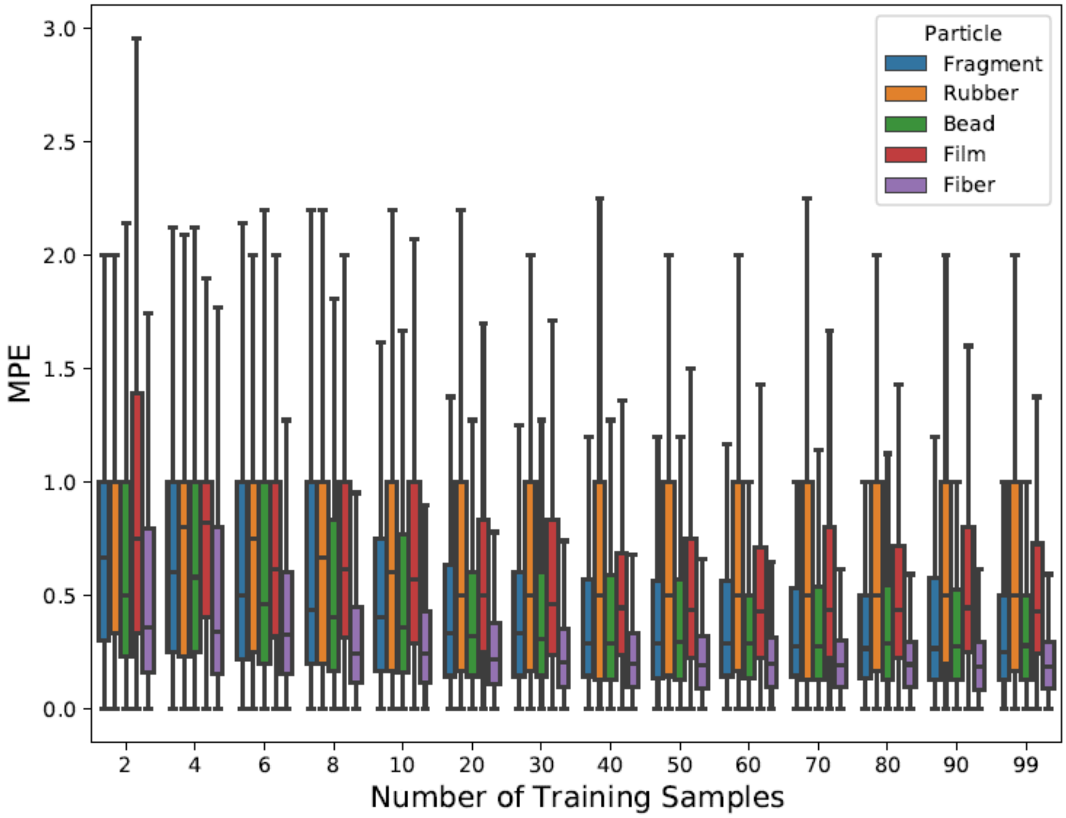


**Figure S13(b): Effect of Training Dataset Size on KRR Prediction Accuracy, Composition 9, Boxplot**

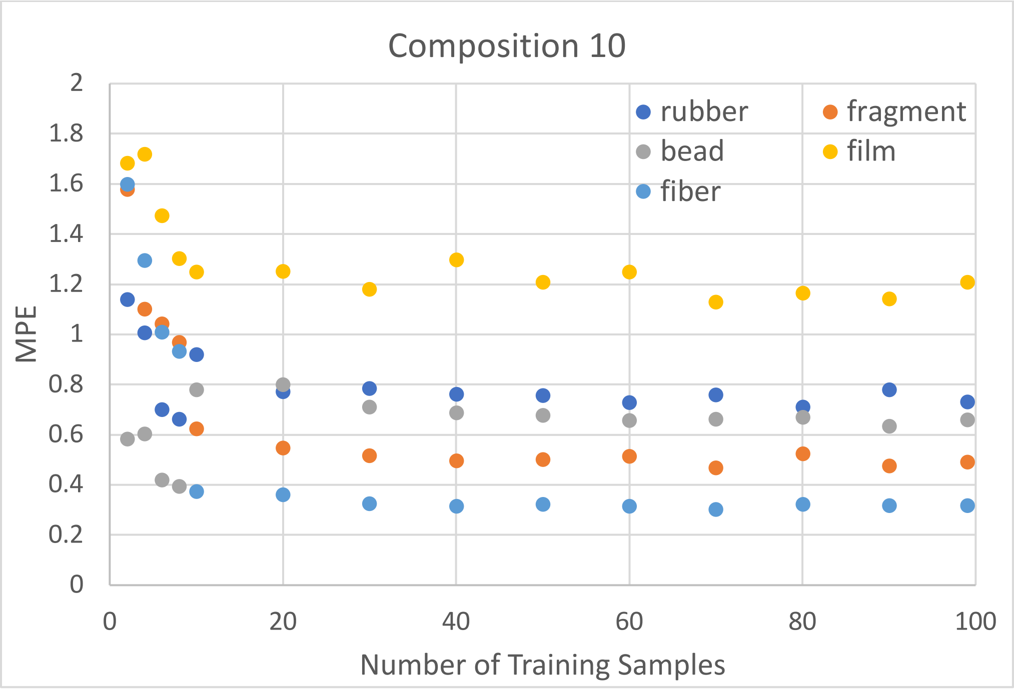


**Figure S14(a): Effect of Training Dataset Size on KRR Prediction Accuracy, Composition 10, Average**

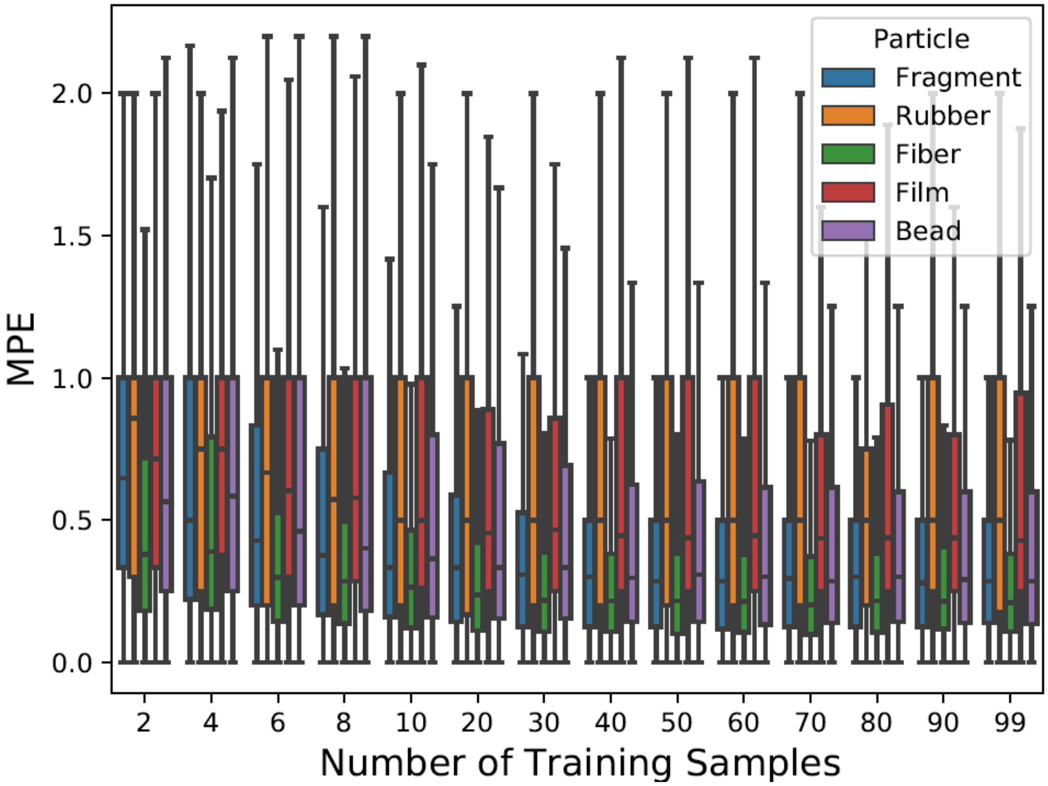


**Figure S14(b): Effect of Training Dataset Size on KRR Prediction Accuracy, Composition 10, Boxplot**

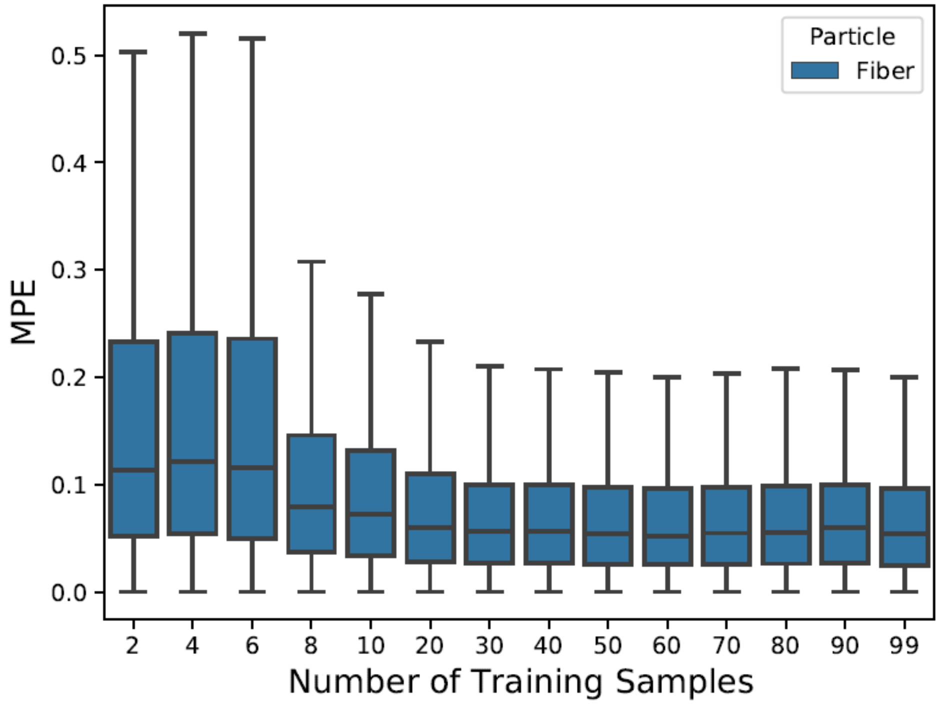


**Figure S15: Effect of Training Dataset Size on KRR Prediction Accuracy, All Fiber, Boxplot**

**Section S5: Performance on Real Samples**

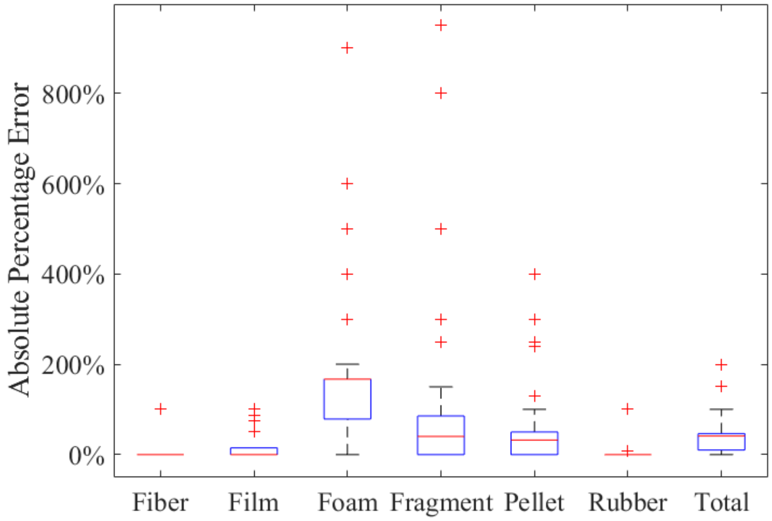


**Figure S16: Mean Percentage Prediction Error of Kernel Ridge Regression on the Great Lake Dataset**

**Section S6: Implementation Examples**

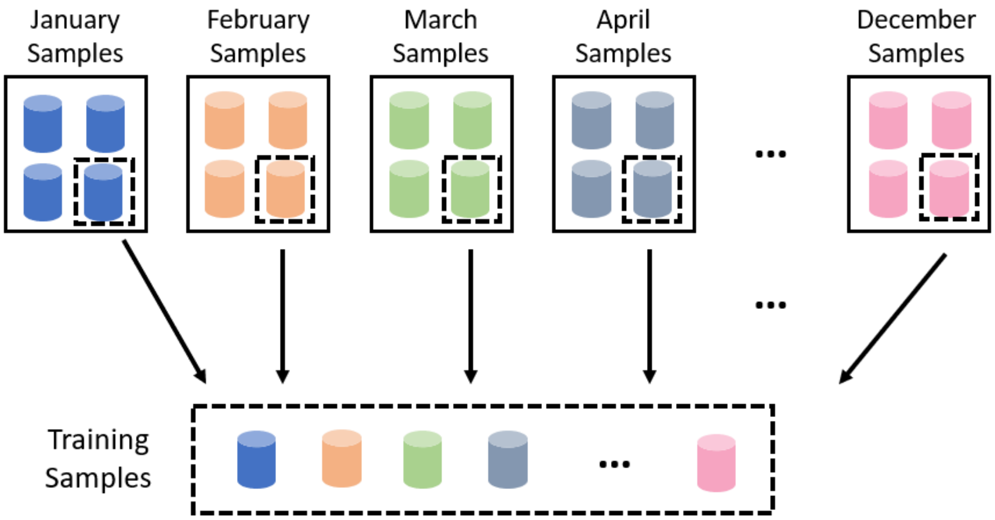


**Figure S17: Temporal Application: part of the samples collected from each time period are used to form a training dataset, train the model, and predict the rest using the model.**

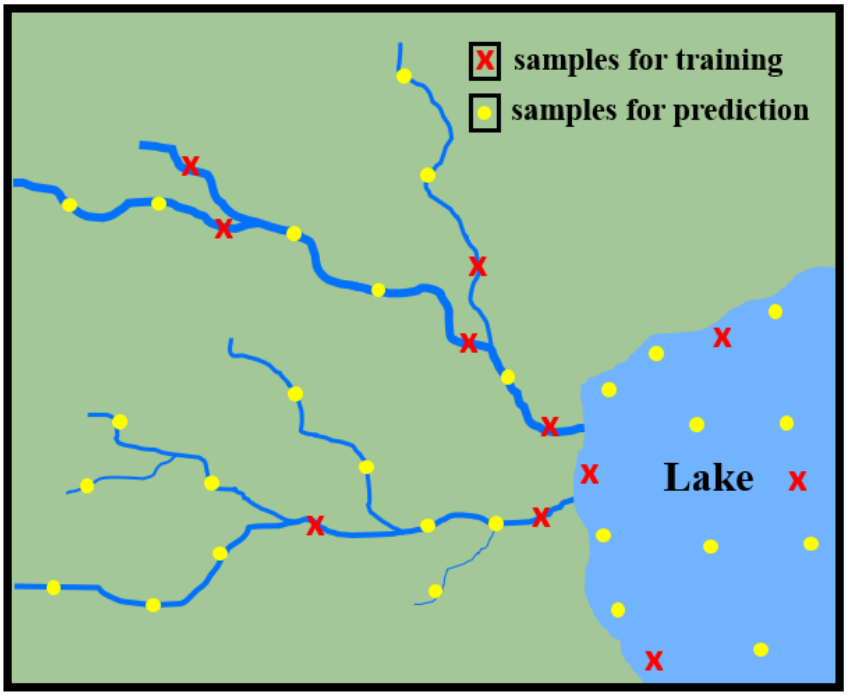


**Figure S18: Spatial Application: samples are collected from many sites, and some samples are used to form a training dataset, train the model, and predict the rest using the model. The red crosses represent samples that will undergo both manual counting and weight measurement; whereas, the yellow dots represent samples that will only be weighed and whose microplastic count will be predicted from the trained KRR model.**

**Table S5: Homogeneous Sample Application: for samples with homogeneous composition, some samples (denoted as training) are chosen to form the training dataset. They will be manually counted and weighted. The rest of samples (denoted as predicting) will only be weighted, and their microplastic count will be predicted from the trained KRR model.**

| Sample | 1 | 2 | 3 | 4 | 5 | 6 | 7 | 8 | 9 | 10 | 11 |
|---|---|---|---|---|---|---|---|---|---|---|---|
| Weight (g) | 0.134 | 0.148 | 0.548 | 0.625 | 0.725 | 0.847 | 1.24 | 1.96 | 2.05 | 3.15 | 4.87 |
| Training | √ | | | | √ | | | √ | | | √ |
| Predicting | | √ | √ | √ | | √ | √ | | √ | √ | |
